# Designing AI-programmable therapeutics with the EDEN family of foundation models

**DOI:** 10.64898/2026.01.12.699009

**Authors:** Geraldene Munsamy, Gavin Ayres, Carla Greco, Keith Kam, Gus Minto-Cowcher, John St. John, Tanggis Bohnuud, Matthew Bakalar, William Chow, Robert Pecoraro, Marcelo D.T. Torres, Aaron Kollasch, Marcus Leung, Hassan Sirelkhatim, Francesco Farina, Connor McGinnis, Srijani Sridhar, Daniel Anderson, Francesco Oteri, Ali Taghibakhshi, Jeremie Dona, Tyler Shimko, Cedric Steenbeke, Alexandros Papadopoulos, Malcolm Krolick, Fabian Spoendlin, Purba Gupta, Sandeep Kumar, Anne Bara, Jared Wilbur, Noelia Ferruz, Timur Rvachov, Fangping Wan, Hanqun Cao, Hyun-Su Lee, Japan Mehta, Raphael Chaleil, Valerio Pereno, Sid Potti, Chris Emerson, Roy Tal Dew, Kevin K Yang, Eric Nguyen, Neha Tadimeti, Jillian F. Banfield, Alicia Frame, Emma Bolton, David Ruau, Rory Kelleher, Anthony Costa, Kimberley Powell, Cesar de la Fuente-Nunez, Glen-Oliver Gowers, Oliver Vince, Jonathan Finn, Philipp Lorenz

**Affiliations:** Basecamp Research, University of Oxford; NVIDIA Corporation, University of Oxford; University of Pennsylvania, University of Oxford; Johns Hopkins University, University of Oxford; Department of Statistics, University of Oxford; Centre for Genomic Regulation, Barcelona, Microsoft Corporation; Stanford University, Microsoft Corporation; Microsoft Research, Microsoft Corporation; University of California, Berkeley, Microsoft Corporation; CoreAI, Microsoft Corporation

## Abstract

The ability to interpret, modify, and design DNA has driven many of the most significant advances in modern medicine, from diagnostics, biologics, and vaccines to cell and gene therapies. However, the inherent complexity of biological systems means that most modern medicines are still engineered using bespoke, labor-intensive processes.

To address the need for a generalisable and programmable approach to therapeutic design, we introduce the EDEN (environmentally-derived evolutionary network) family of metagenomic foundation models, including a 28 billion parameter model trained on 9.7 trillion nucleotide tokens from BaseData^1^. This dataset, at the time of training, contained more than 10 billion novel genes from over 1 million new species, and is intentionally enriched for environmental and host-associated metagenomes, phage sequences, and mobile genetic elements, enabling the model to learn from diverse and novel cross-species evolutionary mechanisms and apply them to key challenges in human health.

EDEN achieves state-of-the-art performance across a series of predictive and generative genomic and protein benchmarks. To demonstrate the models’ broad applicability across biology, we evaluate EDEN’s capacity for programmable therapeutic design by challenging a single architecture to design biological novelty across three distinct therapeutic modalities, disease areas and biological scales:

(i) large gene insertion, (ii) antibiotic peptide design, and (iii) microbiome design.

First, we demonstrate AI-programmable Gene Insertion (aiPGI), in which EDEN designs *de novo* large serine recombinases (LSRs) capable of inserting large pieces of DNA at desired target sites in the human genome when prompted only on 30 nucleotides of DNA sequence from the desired target site. In low-N experimental validation, EDEN generated multiple active recombinases for all tested disease-associated genomic *loci* (ATM, DMD, F9, FANCC, GALC, IDS, P4HA1, PHEX, RYR2, USH2A) and 4 potential safe harbor sites in the human genome. EDEN achieves an overall functional hit rate of 63.2% across diverse DNA prompts when prompted on only 30bp of DNA from outside the training data.

50% of EDEN-generated LSRs were active in human cells, achieving therapeutically relevant levels of CAR insertion in primary human T cells. We also show that EDEN can generate active bridge recombinases when prompted on the associated guide RNA alone, with sequence identities to training and public data as low as 65%. These results pave the way for a new generation of cell and gene therapies by opening the door to rapid, programmable and site-specific integration of large genetic payloads without double-strand breaks. This offers an alternative to the safety, efficiency and payload limitations inherent in viral or nuclease-based editing at thousands of currently intractable human therapeutic targets.

Second, we use the same model to generate a focused low-N library of novel antimicrobial peptides where 97% showed activity, with top candidates achieving single-digit micromolar potency against critical-priority multidrug-resistant pathogens.

Third, to demonstrate that EDEN captures *inter*-genomic features, we design a gigabase-scale microbiome with over 94,000 synthetic metagenomic assemblies, including prophage genomes and correct cross-species metabolic pathway completions. The EDEN-generated synthetic microbiome covers 9,067 species with a biome-specific taxonomic accuracy of 99%. Over 1,500 of the generated species were outside the fine-tuning dataset while retaining the correct microecological properties and biome association, thus significantly expanding genetic and taxonomic diversity.

Together, these results establish a new strategic direction for AI-programmable therapeutics, in which a single foundation model architecture designs candidate therapeutics across diverse modalities and disease areas. This suggests that the combination of billions of years of evolutionary data with specific therapeutic records offers a clear, scaling-driven path to making therapeutic design a predictable engineering discipline.

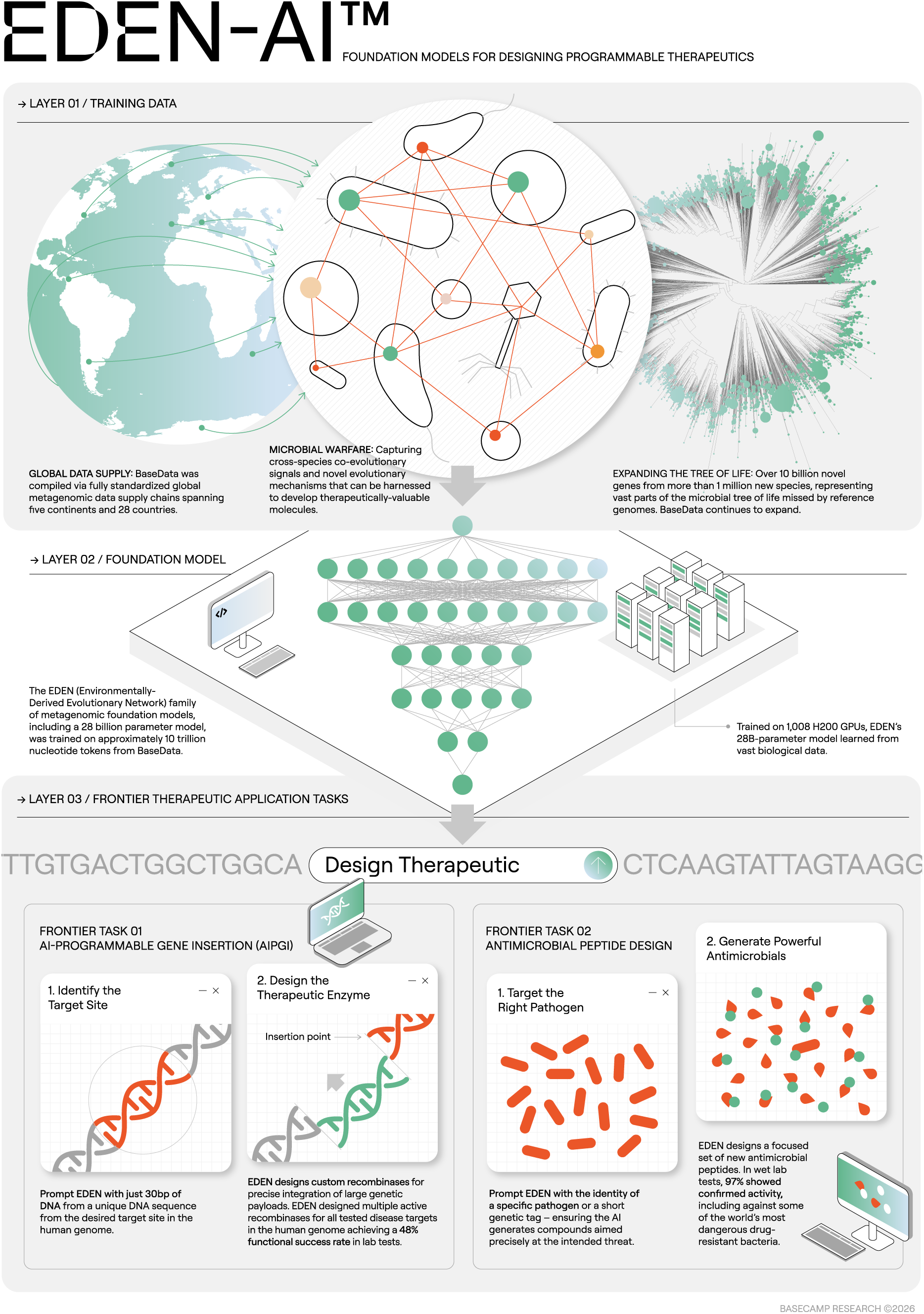

## Introduction

### Background & motivation

The ability to generalisably and predictably design therapeutics holds the potential to transform human health, offering the ability to address complex pathologies with speed, efficacy and precision. However, despite decades of progress in genomic engineering and the foundational premise of systems biology that biological systems are non-random, logical and inherently computable, the systematic programming of biological outcomes has so far remained largely intractable. Moving therapeutic design towards a deterministic engineering discipline represents a major, and still largely unrealized, opportunity in modern medicine.

This paper frames the current intractability of biological programming as an engineering bottleneck arising from the profound information asymmetry between the complexity of biological systems^2,3^, and two interlinked technical deficits: (a) the scarcity of diverse primary biological information and (b) the lack of computational systems capable of translating this information into medicine^4,5^. These deficits arise from the limitations of physical laboratories as the primary tool for large scale data collection and human cognition as the primary engine for data interpretation; neither of which has the capacity to match the scale or complexity resulting from four billion years of evolution.

In previous work^1^, the authors presented a new approach for surmounting the first deficit; a new, globally scaled and partnership based supply chain of evolutionary genomic data. The resulting dataset, BaseData, contained nearly 10 billion genes from over 1 million new species at the time of publication, and is capable of growing at over 2 billion genes per month.

AI models offer a promising new paradigm for surmounting the latter deficit; by learning statistical patterns within large biological datasets, generative foundation models are unlocking increasingly sophisticated and generalisable biological design capabilities – generating functional proteins, regulatory elements and even complete genomes in response to complex queries, often without the need for exhaustive experimental screening^6–13^. However, as detailed in our previous work^1^, the full potential of these models for biological design could likely be enhanced significantly by leveraging additional diverse biological sequence data, especially those derived from the complex interplay between hosts, mobile genetic elements and their environments.

On this basis, we hypothesize that progress towards true programmable biology will require expanding the training datasets of generative models to include increasingly large quantities of diverse evolutionary data, far beyond the constraints of current publicly available resources. If this hypothesis is true, we would expect these models to learn increasingly universal design principles from this data and progressively improve the predictability, accuracy, and controllability of the computational design of biological code.

To test this hypothesis, we introduce the EDEN (Environmentally-Derived Evolutionary Network) family of foundation models, the largest of which was trained on 9.7 trillion evolutionary nucleotide tokens from BaseData^1^, with no human, lab or clinical data in the pre-training dataset. We evaluate EDEN’s capacity for programmable therapeutic design by challenging a single architecture to design biological novelty across three distinct therapeutic modalities, disease areas and biological scales: (i) large gene insertion, (ii) antibiotic peptide design, and (iii) microbiome design.

Through this, we demonstrate that a single foundation model, learning from a greater diversity of genomic evolution than previously available, can drive the design of novel potential medicines in response to a range of therapeutically-relevant queries. Together, these results open the path and indicate a route towards unified AI systems capable of designing therapeutic candidates across diverse disease areas and modalities.

### The evolution of biological design

The treatment of biology as a designable system represents the culmination of a century-long shift from viewing life as a “vital force” to seeing it as a modular information technology. This trajectory began with Darwin and Mendel, who established the optimization logic (natural selection) and the discrete units of inheritance (genes) that allow life to operate as a set of defined, logical systems^14,15^. The theoretical convergence of physics and information theory saw Schrödinger predict the molecular storage for this code, while Shannon, Turing and von Neumann provided the mathematical framework that formalized biological self-replication as a computable process^16–18^.

In parallel to these theoretical insights, our rapidly improving ability to physically read, write and design DNA has driven decades of progress in medicine and biotechnology^15,19,20^. Recombinant DNA technology now underpins the production of therapeutic proteins, including hormones, peptides, antibiotics, and antibodies used to treat diseases such as diabetes, cancer, bacterial infections and autoimmune disorders^21–24^.

More recently, the field has moved beyond simple protein production towards the editing and engineering of more complex multi-component systems: for example, genome editing technologies like CRISPR/Cas9 have moved from research to clinic within a decade^25,26^. Already, CRISPR-based therapies are curing genetic diseases in clinical trials (for example, *ex vivo* edited hematopoietic cells for β-thalassemia and sickle cell disease) and have recently proven effective in a rapidly developed personalized therapy for an infant with CPS1 deficiency, an ultra-rare genetic metabolic disease^27,28^

Moving up the ladder of complexity, engineered cell therapies such as CAR-T cells – T-cells genetically modified with synthetic receptors – have achieved unprecedented success in refractory leukemias, with seven therapies approved by early 2026^29,30^. Simultaneously, the maturation of mRNA platforms has introduced a high-velocity modality, enabling the “programming” of personalized cancer vaccines that translate patient-specific tumor mutations directly into therapeutic instructions^31,32^.

However, despite these breakthroughs, biological engineering remains a comparatively artisanal endeavor when compared to true systematic engineering disciplines. It is still common for complex therapeutic development to depend on labor-intensive, stochastic screening campaigns^33,34^. This bespoke, trial-and-error approach is (a) functionally unscalable, (b) largely restricted to “low-hanging fruit” targets where natural proteins can be easily repurposed, and (c) fundamentally incapable of addressing the vast majority of complex polygenic or multi-factoral pathologies. Consequently, the immense therapeutic potential of programmable biology remains largely latent, constrained by our inability to design *de novo* function.

The hypothesis within this paper is that the current constraints in therapeutic design arise from a profound disparity between the information content within biological systems and the processing bandwidth of our engineering tools. Evolution operates on a timescale of eons across a planetary-scale ‘laboratory’, optimizing fitness within an extremely high-dimensional combinatorial landscape that dwarfs laboratory capacity and human cognition.

Physical laboratories, while essential for validation, act as comparatively low-throughput filters for data collection. Even the most advanced high-throughput screening campaigns capture only an infinitesimal fraction of the theoretical sequence space, leaving the vast majority of the “design universe” unexplored. The authors have discussed this extensively in previous work^1^. Compounding this data sparsity is a cognitive limit: the rules governing biological function, defined by high-order epistasis, long-range interactions, and environmental context, are too subtle and multidimensional for human intuition alone to decipher.

Consequently, therapeutic development stays in a cycle of iterative screening. As this laboratory-intensive approach scales linearly, it is fundamentally incapable of matching the exponential complexity of disease biology. This bottleneck necessitates a need for generalisable and programmable biological design – algorithms or computational systems that can learn evolutionary grammar at a scale sufficient for designing novel therapeutic constructs on demand across diverse modalities and disease areas, in response to specific, and, ultimately, personalized therapeutic queries.

### Foundation models in biology

In parallel to the advancements in genomic engineering, foundation models - large-scale deep learning models trained on broad, multi-billion to trillion-token scale datasets - offer the field a way past the cognitive bottlenecks outlined above. These models, trained on massive and heterogeneous datasets, often using self-supervised objectives, exhibit the capacity to generalize across a wide range of downstream tasks without task-specific retraining^35^ and have revolutionized domains including natural language, computer vision and robotics.

As model architectures matured - with the introduction of the Transformer in 2017 – foundation models advanced further through rapid growth in available compute and the assembly of large-scale datasets^35–37^. Together, efficient architectures, abundant data, and increased computational resources enabled more systematic study of how empirical performance varies with model and dataset scale. These studies, often referred to as scaling laws, show that, across broad regimes, performance improvements are well approximated by power-law relationships in model size, dataset size, and training compute, thereby reframing model progress as an engineering problem of allocating capacity, data, and computation effectively^38–41^.

Empirical studies demonstrated that training models on increasingly large datasets produced emergent behaviors such as in-context learning and compositional reasoning^42^. This principle guided the development of large language models such as the Llama and GPT families of large language model^43,44^, trained on datasets of hundreds of billions to trillions of tokens derived from diverse web, book, and code sources. These models have had a profound societal impact, transforming how we interact with machines and enabling breakthroughs in software development, education and healthcare, through enhanced personalisation and automation^45–47^.

In turn, the application of large AI models to biological problems has transformed computational biology and our understanding of molecular design and properties, with use cases spanning industrial processes, synthetic biology, and therapeutic applications^48^. This is exemplified by AlphaFold2, which, when trained on multiple sequence alignments, 3-dimensional protein structures, and atomic-coordinate supervision, achieved near-experimental accuracy in protein structure prediction, work that was ultimately awarded the 2024 Nobel Prize in Chemistry^49–51^. Building on this, AlphaFold3 expands training to protein–protein, protein–nucleic-acid, and protein–ligand complexes and uses diffusion-inspired refinement to improve interaction and assembly accuracy, while other biomolecular foundation models such as those from the Chai or Boltz family combine generative, diffusion-based, and graph-neural approaches to enable joint sequence–structure generation and functional binding design^52–59^.

In addition to this, language modeling has been an additional major driving force in biological foundation modeling, especially in protein and DNA design. Protein language models treat amino-acid sequences as unlabeled data for self-supervised learning, supporting downstream tasks such as supervised fitness prediction, domain annotation, mutation effect prediction, structure prediction, and sequence generation. Early autoregressive models trained on UniProt/UniRef, including ProGen and ProtGPT2, showed that de novo samples can preserve natural amino-acid propensities^8,10–12,60^. ZymCTRL added Enzyme Commission label conditioning to steer sequences toward specific catalytic functions, and PoET then framed protein families as “sequences-of-sequences,” using family-aware transformers to improve zero-shot variant effect prediction and controllable family-conditioned generation^12,61,62^. Diffusion-based models such as EvoDiff and DPLM enabled order-agnostic generation, while ProGen3 and Dayhoff demonstrated that training on metagenomic data increases the quality and diversity of generated sequences^11,63–65^. Meanwhile, ESM models demonstrated that massively scaled self-supervised training on protein sequences yields embeddings capturing structural grammars^61,62,66,67^. However, it remains difficult to precisely steer these models for desired functions, such as DNA-protein interactions or antimicrobial activity, as most models do not consider genomic or community context.

Beyond proteins and molecular interactions, genomic language models have extended to entire genomes by treating the genome as a learnable language whose syntax encodes regulatory function beyond individual Open Reading Frames (ORFs)^68,69^. Encoder-only models such as DNABERT first demonstrated that transformers pretrained on k-merized genomic sequences could learn promoter and splice-site grammars across species^70,71^. Subsequent DNA foundation models expanded training data or context, such as the Nucleotide Transformer enabling zero-shot predictions of regulatory elements or HyenaDNA expanding genomic modelling context to the megabase scale^72,73^. Genome-scale generative models such as Evo and Evo2 further unified molecular-to-genome modeling across long-range genomic interactions, supporting variant effect inference and realistic genome synthesis^6,7^. Impressively, Evo2 has done so by scaling the training data to all known domains of life^7^. AlphaGenome extends this paradigm to multimodal prediction, coupling sequence inputs with chromatin and structural readouts^74^.

Across the various design tasks, whether based on protein-or genomic foundation models, the prediction and design of protein-DNA interactions have proven to be particularly challenging^75^. In particular, at a molecular level, DNA’s primarily sequence-independent double-helix structure complicates accurate modeling and design of protein interactions with specific DNA sequences. Compared to protein-small molecule, or protein-protein interactions, protein-DNA interactions often score worse across the most recent protein-ligand design models^52–56^. In this domain, significant progress has been made, for example with the design of sequence-specific DNA-binding proteins with helix-turn-helix domains using RFDiffusion^76^. However, while various design campaigns based on biological deep learning models have indeed generated functional DNA-binders or-editors, there is room for further improvement and generalisation: their design from biological deep learning models has relied on high-throughput experimentation in the millions of variants range, prompting on the entire protein itself, up to ten epochs of fine-tuning, as well as the restriction to specific subfamilies^6,77,78^.

In this paper, we suggest that heavy dependence of current models on extensive experimental iteration and the inability of current models to reliably execute complex tasks necessary for multi-modality therapeutic design may arise from a fundamental deficit in the scale and diversity of the available training data, whether that be sequence-specific, structural, or otherwise^1,51,79–82^.

Effective modeling of complex therapeutic modalities depends on models’ ability to learn higher-order biological interactions, for example cross-species DNA-protein interactions, host-pathogen interactions, and other multi-species co-evolutionary signals, especially when designing more complex therapeutic modalities such as peptides or cell and gene therapies^83–86^. For example, modeling Cas9 nucleases or Large Serine Recombinases (LSRs) necessitates sourcing immense quantities of high quality, diverse primary data on the proteins and their associated non-coding elements, including guide RNAs and attachment sites, which dictate target specificity^1,77^.

In this context, deep learning models in this space have frequently relied on the usage of reference genomes, often derived from cultured isolates^6,7,73^. Whilst these are valuable resources for microbial genomics, public biological sequence databases frequently fail to capture the natural evolution within microecological complex environments (metagenomes, microbiomes) at scale that give rise to vast, otherwise unmapped parts of the tree of life^87–89^. This data sparsity has consequences for models’ ability to learn the higher-order biological interactions discussed above, and thus is one of the key limits on progress towards true programmable therapeutic design.

### Introducing EDEN: learning from evolution

Returning to our central hypothesis, that a path to programmable biology lies in expanding training data distributions of generative models to capture broader evolutionary context – we face two interlinked technical challenges: the lack of known biological information and the lack of cognitive bandwidth necessary to fully interpret the information that we do have.

From this hypothesis, it follows that, if these deficits could be systematically addressed, an improvement in the predictability, accuracy, and controllability of computational biological design would be observed.

To address the first of these challenges, the authors previously published BaseData^1^ which has increased the known non-redundant sequence diversity both globally and across key gene editing protein families of interest, contextualized with over 4 times the genomic context and over a million new species compared to comparable public biological sequence databases^1,11,81,82^.

Now, to address the second of these challenges, we introduce the EDEN (Environmentally-Derived Evolutionary Network) family of foundation models trained on up to 9.7 trillion nucleotide tokens from BaseData. EDEN uses a Llama3-style architecture, scaling from 100 million to 28 billion parameters. Trained on up to 1.95×10^24^ FLOPs, the EDEN models are some of the largest foundation models ever trained and achieve state-of-the-art performance across a range of predictive and generative genomic and protein benchmarks.

To evaluate EDEN’s ability to learn from evolution to predictably, accurately, and controllably design therapeutic modalities, we moved beyond traditional benchmarks to test the models on three distinct design challenges that span a range of modalities, disease areas and biological scales: (a) large gene insertion, (b) peptide design and (c) synthetic microbiomes.

In the next section, we discuss the background and motivation of each of these therapeutic tests, establishing the existing benchmarks and the technical and medical implications of these achievements.

### EDEN designs therapeutics across scales, modalities, and disease areas

#### AI-Programmable Gene Insertion (aiPGI)

Despite major advances in genome engineering, current technologies still fall short of delivering programmable, scalable solutions for repairing or rewriting the human genome. CRISPR nucleases, base editors, and reverse-transcriptase-based editors have enabled targeted correction of single-nucleotide changes or short indels, but these strategies remain inherently mutation-specific^26,90^. In practice, this means a therapeutic edit must be uniquely designed for the exact pathogenic variant a patient carries. Even within a single disease, different patients often carry distinct mutations, and in many disorders the pathogenic landscape spans hundreds to thousands of allelic variants^91–93^. Consequently, CRISPR-based correction scales poorly: each patient, or small patient subgroup, requires its own bespoke edit, limiting both clinical generalisability and the feasibility of broad therapeutic deployment^93^.

Programmable Gene Insertion (PGI), the ability to efficiently insert large pieces of DNA into specific genomic locations, is a significant challenge for the gene editing field. PGI has the potential to address many limitations of current gene therapy and gene editing systems, such as treating very heterogenous genetic diseases with a single therapy, insertional oncogenesis (due to random integration), non-native gene expression (using safe harbor sites), and loss of episomal expression over time (particularly in pediatric patients). In addition to increasing the safety and efficacy of current cell therapy approaches (e.g. CAR-T engineering), PGI enables the insertion of healthy copies of genes into their correct endogenous locations in patients with genetic disease. This would enable native regulation, (avoiding over or under-expression), a single product that will treat most if not all patients (mutation agnostic), and allow for a one-time cure that grows with the patient.

Large Serine Recombinases (LSRs) are an abundant class of enzymes found in nature that have many properties that make them attractive for PGI applications. They are very small, can efficiently integrate large pieces of DNA (>30 kb^94^), have a predictable integration profile^95^, and are not reliant on DNA damage or host DNA repair pathways, meaning they are efficient in both dividing and quiescent cells.

The key challenge with developing recombinases for human therapeutics is that each natural LSR has evolved to integrate into a unique bacterial DNA sequence, none of which are at sites relevant for human therapeutics. While there have been efforts to re-direct LSRs to novel sites^96–98^, most efforts use laborious wet-lab evolution and/or involve appending additional domains to the LSR, which can have a negative effect on both efficiency and size. LSR-mediated PGI has indeed shown to be feasible in a non-human primate model^99^, however this approach was limited by complexity and manufacturing challenges since it required a Cas9 nickase and reverse transcriptase in order to make LSR-mediated insertion programmable.

An elegant solution would be to be able to design programmability directly into the recombinase itself. However, addressing this inverse design problem - mapping a desired DNA target back to a functional protein sequence - requires a generative model that understands the high-dimensional evolutionary logic coupling LSR amino acid sequences to their specific DNA targets.

To validate EDEN’s capacity for AI-Programmable Gene Insertion (aiPGI), we use EDEN to generate *de novo* large serine recombinase proteins when prompted with only the desired genomic target site. Recombinase design serves as a rigorous benchmark for programmable biology by testing the model’s capacity for writing complex biological instructions from a comparatively small prompt, whilst simultaneously optimizing DNA-binding specificity and catalytic efficiency.

We show that, when prompted with only 30 nucleotides of DNA representing the desired attB genomic target site, EDEN generates multiple active recombinase proteins for all tested disease-associated human genomic *loci* (ATM, DMD, F9, FANCC, GALC, IDS, P4HA1, PHEX, RYR2, USH2A) and four potential safe harbor sites in the human genome. Over all prompts, EDEN achieves a functional hit rate of 53.6%. Top tier variants exhibit biochemical activity on par with any natural recombinases screened to date and several high-performing candidates shared as little as 52% sequence identity with the parental protein, indicating that EDEN is learning biological ‘grammar’ and efficiently navigating the vast evolutionary sequence landscape.

In parallel to attachment-site-prompted LSR design, we showcase EDEN’s ability to address aiPGI in an orthogonal approach by designing active bridge recombinases (BRs) which have recently been studied and developed as an RNA-programmable alternative to LSRs (Durrant et al. 2024; Perry et al. 2025). EDEN-generated BRs were prompted solely on their corresponding non-coding RNA sequence and exhibit sequence identity to the training and any public BRs as low as 65%.

By enabling the site-specific integration of large genetic payloads in a single protein without the genotoxicity associated with double-strand breaks and without the requirement for a guide RNA, these results suggest that EDEN has the potential to systematically address the complexity, safety and payload limitations of current viral and nuclease-based editing in gene therapy, paving the way for new generations of complex and curative cell and gene therapies to treat a much broader range of indications than is currently accessible.

While these *de novo* designs are potent functional hits, it is important to acknowledge that they will require downstream optimization before becoming clinic-ready medicines. Nonetheless, as the EDEN models continue to improve, this capability opens the door to a programmable toolkit for safe, large-payload gene integration and is a powerful proof point on the route to designing personalized therapeutics within the complex requirements of clinical intervention.

### AI-based antimicrobial peptide design

The escalation of antimicrobial resistance (AMR) has created an urgent imperative for new therapeutics, as drug-resistant “superbugs” are now recognized among the World Health Organization’s top global health threats^100–103^. In particular, critical-priority multidrug-resistant pathogens – exemplified by the ESKAPE bacteria (e.g. carbapenem-resistant *Acinetobacter baumannii*) – pose imminent dangers that could usher in a post-antibiotic era without effective countermeasures^104,105^. Motivated by recent findings that AI can be used to accelerate antibiotic discovery^106^, and that antimicrobial peptides (AMP) can be discovered from biology, including within microbiomes^86,107^, the human proteome^108^, and ancient biology^109–111^, we envisioned using EDEN for the generative design thereof.

Biologically, AMPs constitute a diverse class of short peptides produced by a wide range of organisms, as well as identified in numerous environmental microbiomes, and many display broad-spectrum activity through mechanisms that are less prone to conventional resistance^22,84^. Yet AMP discovery and design, whilst showing successes both from existing genomic resources as well as through machine-learning methods^110,112^, could be scaled significantly by widening the phylogenetic and environmental diversity beyond what is represented in public resources, which are otherwise limiting the ability of machine learning models to generalize fully across the antimicrobial sequence landscape towards programmably targeting the most relevant pathogens.

To validate EDEN’s ability to design functional therapeutic candidates within near-infinite sequence spaces, we applied the model to the *de novo* design of AMPs. Experimental validation revealed that 32 of the 33 EDEN designed peptides (97%) were functional, demonstrating high potency against WHO critical-priority pathogens, including multidrug-resistant *Acinetobacter baumannii*^100,101^. By achieving such high precision without iterative experimental cycles, we show the potential of an AI-driven framework for responding to the global antimicrobial resistance crisis. While these *de novo* peptides exhibit potent activity, they remain early-stage candidates requiring further optimization for stability, toxicity, and pharmacokinetic properties before clinical application.

### Synthetic microbiome

The ultimate frontier in programmable biology lies in understanding complex multi-species biological systems such as microbiomes, and developing the ability to programmably generate them. Their design requires accounting for emergent properties including metabolic cross-feeding, niche competition, and trophic stability that are absent at the single-genome level^113–117^. One example for such multi-species systems include host-associated microbiomes which are well-established in their role in human metabolic health and carcinogenesis^118–121^, with precision microbiome editing having recently shown promise for human health and disease applications^122^. Modeling this higher-order logic represents a distinct challenge from molecular design, requiring the internalization of ecological rather than just structural syntax.

While previous generative biological foundation models have achieved success in designing open-reading frames, mobile genetic elements, and even whole genomes, they have largely stopped at the organismal boundary^7,123^. Current approaches fail to capture the “dark matter” of interactions that dictate community survival. Without modeling the cross-species dependencies inherent in natural environments, the *de novo* design of a stable, functional microbiome remains out of reach for models trained on isolated reference genomes.

To validate EDEN’s capacity for design beyond the individual genome, we challenged the model to generate a fully synthetic, gigabase-scale microbiome. Leveraging the cross-species evolutionary information inherent in its metagenomic training, EDEN generated a synthetic host-associated community containing phage genomes and biome-specific metabolic pathway completions across different synthetic assemblies. 99% of generated species had the correct biome association. By generating a microbiome that coheres at the gigabase scale, we show that EDEN captures statistical regularities at the metagenome level. While these synthetic microbiomes require experimental instantiation and validation, the *in silico* results suggest the feasibility of moving from the design of individual molecules or genomes to a larger scale at the biological community level.

### Toward a unified model for programmable therapeutics

By validating the EDEN models across three different biological scales, disease areas and therapeutic modalities, we demonstrate that a single foundation model, learning from a higher diversity of genomic evolution than previous models, can drive towards more predictable engineering of novel potential therapeutic candidates in response to sophisticated and specific therapeutically-relevant queries.

Together, these results indicate that training on more evolutionary data is an important and likely underappreciated part of the path towards unified AI systems capable of designing therapeutic candidates across multiple diverse disease areas and modalities. If this trajectory continues, these systems hold the potential to evolve from predicting properties to designing full curative therapeutic interventions, bringing currently intractable pathologies within the reach of personalized and programmable medicine.

## Results

### Training the EDEN model family

EDEN was trained on BaseData, a training data corpus enriched for environmental and host-associated metagenomes and purpose-built for foundation model training^1^. We have previously shown how BaseData expands the known sequence space compared to several public resources, and include the comparison between BaseData and the metagenomic portion of OpenGenome-2 (OG2), the dataset used for training the frontier Evo2 genomic foundation model^7^ (Figure 1A). The range of environmental features BaseData has been sourced from is also shown, representing the genomic sequence space as it relates to the sample biome (Figure 1B) and pH (Figure 1C). This type of metadata is largely not consistently captured in public resources. Beyond the sequence space itself, we show the distribution of contig nucleotide length and ORF count (Figure 1D-E), with BaseData assemblies showing significantly larger genomic context compared to those from the metagenomic portion of OG2.

**Figure 1:**
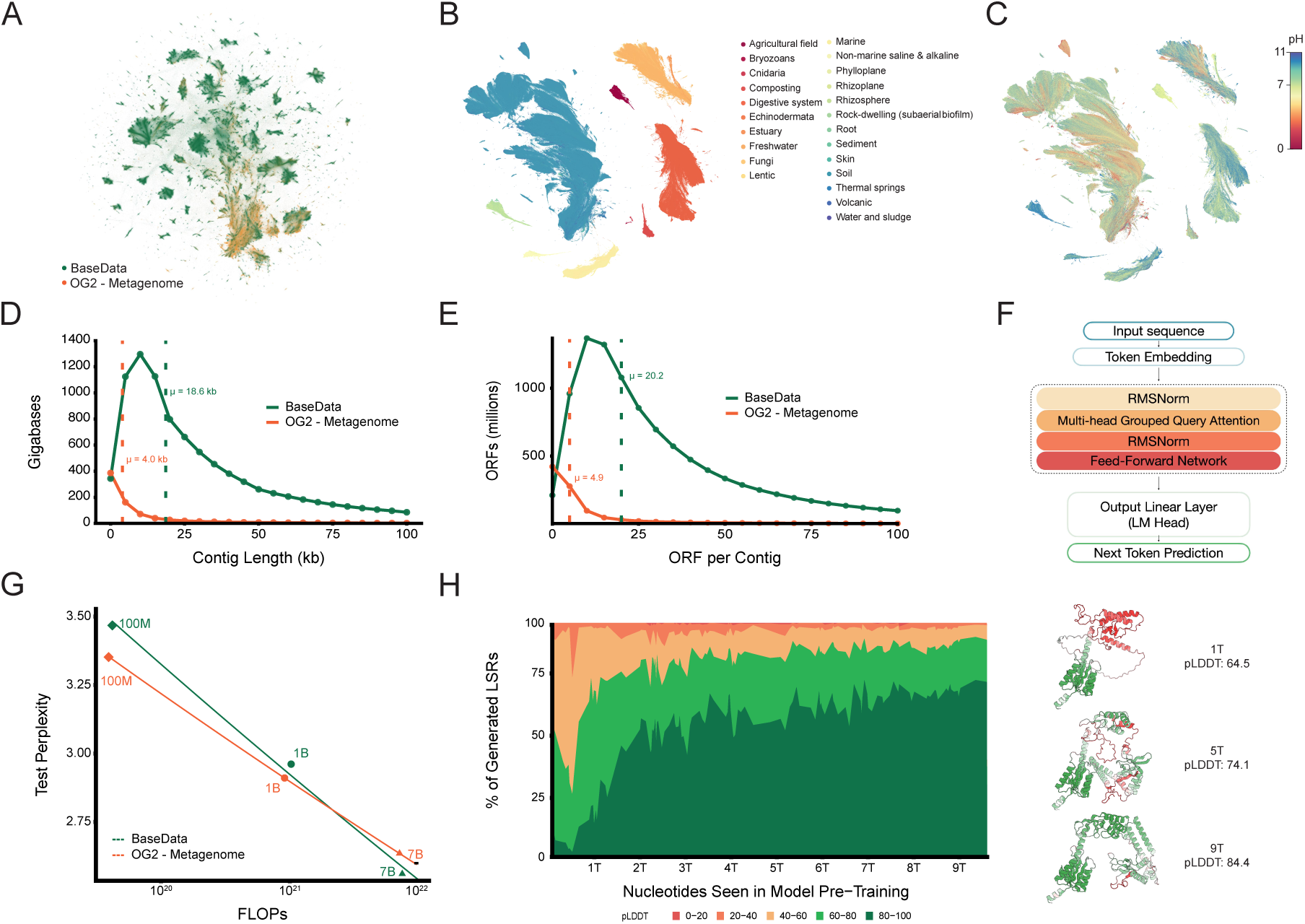
The EDEN model family. **A** UMAP of metagenomic assemblies from BaseData and the metagenomic portion of OpenGenome2. **B** UMAP of metagenomic assemblies from BaseData colored by biome origin. **C** UMAP of metagenomic assemblies from BaseData colored by pH. **D** Distribution of contig lengths across metagenomic databases, showing a median contig length of 18.6 kb for BaseData and 4.0kb for OG2 (metagenomic). **E** Distribution of ORF number per contig across metagenomic databases, showing a median of 4.9 ORFs per assembly in OG2 (metagenomic) and 20.2 in BaseData. **F** Llama3.1-like architecture used for EDEN training. **G** Test perplexity vs FLOPs across the EDEN family of models at 100 million, 1 billion, and 7 billion parameters, utilized as a basis for the decision to scale the EDEN model family to a 28 billion parameter model trained on the entirety of BaseData. **H** Distribution of EDEN-28B-generated large serine recombinase (LSR) pLDDT when prompted with 30% of the 5” end of the ORF across the pretraining course. On the right we show example structures of EDEN-generated LSRs from various points (token counts 1 trillion, 5 trillion, and 9 trillion) across the pretraining run.

The EDEN family of models was trained on BaseData using a next-token prediction objective and a context length of 8192 tokens using a Llama3.1-style architecture^1,7,43,124^ (Figure 1F, Methods). Quality-aware scaling laws extrapolate how much computation is required to achieve a desired performance threshold^40^. These scaling laws model the test loss as a function of model size, data volume, and an effective data-quality parameter, with higher-quality data increasing the useful information per token and thereby reducing the computational resources required to reach a performance target^40^.

To explore how different metagenomic datasets influence model performance during scaling, we trained three pairs of EDEN models with 100M, 1B, and 7B parameters on a randomly sampled subset of contigs covering 350 billion nucleotides from the metagenomic portion of OpenGenome2 (OG2; mean contig length ≈ 4 kbp) and BaseData (mean contig length ≈ 18 kbp). To ensure a fair comparison, all models were matched by the number of non-padding nucleotide tokens, and padding-related compute overhead was explicitly corrected when constructing the FLOPs axis in Figure 1G (Methods). Fitting a power law between training FLOPs and test perplexity shows that perplexity decreases more rapidly with compute on BaseData. (Figure 1G).

Consistent with this, while the 100M parameter model trained on OG2 performs better than the 100M model trained on BaseData, the 7B model trained on BaseData achieves lower test perplexity than its OG2 counterpart. This crossover supports a quality-aware interpretation: small models underfit the longer-range structure in the longer metagenomic assemblies in BaseData, while larger models have sufficient capacity to exploit it, extracting more useful information per token^41^. We therefore hypothesize that the observed difference in scaling behavior reflects intrinsic information structure: OG2 seldom presents contiguous genomic context beyond ∼4 kbp, whereas BaseData routinely provides multi-kilobase neighborhoods within a single window. This richer long-range context offers a plausible explanation for the steeper scaling exponent observed for BaseData, consistent with frameworks in which data quality and long-range information content modulate compute efficiency^125^.

This observation also motivates a curriculum-learning strategy. Recent work on continual pretraining suggests that training first with shorter context windows and then transitioning to longer contexts can achieve similar final performance at lower computational cost, consistent with rapid adaptation once long-range dependencies are introduced^126^. In our setting, a short-to-long curriculum could further improve BaseData’s compute efficiency by learning local sequence regularities during an inexpensive short-context phase, then allocating long-context compute to the multi-kilobase genomic neighborhood signal that BaseData frequently contains but OG2 rarely provides.

Motivated by the scaling trends observed (Figure 1G), we trained a 28 billion parameter model, EDEN-28B, on the entirety of BaseData (9.7 trillion nucleotide tokens at the time of training). EDEN-28B attains the lowest test perplexity and falls near the extrapolation of the scaling fit from the smaller models, indicating that EDEN continues to scale efficiently and predictably when trained on the entire dataset (Supplementary Figure 1). In addition to tracking pre-training loss, we periodically evaluated EDEN-28B on biologically relevant downstream tasks, including semantic mining autocompletion and LSR generation. During pre-training, we saved model checkpoints every 1250 steps and, at each checkpoint, generated large serine recombinase (LSR) proteins and assessed them using an *in silico* evaluation pipeline (Methods). We show that the proportion of generated LSR sequences with high pLDDT values increases monotonically across the pretraining run up to the final 9.7 trillion token point (Figure 1H-I). This proportion increases steadily over the course of pre-training, indicating that, as the model is exposed to more data, optimisation progress is accompanied by consistent improvements in a task-aligned confidence metric for the generated proteins.

### EDEN model evaluations

We investigated EDEN’s zero-shot performance on a range of biologically relevant predictive and generative tasks. First, we tested the ability of EDEN to predict mutational effects on protein-coding gene function leveraging deep mutational scanning (DMS) studies, a method commonly used by protein language models and more recently DNA language models^127,128^. DMS involves generating a comprehensive library of sequence variants and experimentally assessing how each mutation influences one or more fitness readouts that reflect the functional performance of the molecule. The likelihood or pseudolikelihood computed by a language model for a DNA or protein sequence can be used to predict its experimental fitness.

EDEN displays state-of-the art performance across genomic and RNA foundation models for this benchmark when averaging across prokaryotic protein-coding genes (Figure 2A). In particular, EDEN achieves higher performance than all other models on β-lactamase *E. coli*, even though this task heavily overlaps with the training data of existing DNA and protein language models, whereas EDEN is trained solely on diverse environmental sequences.

**Figure 2:**
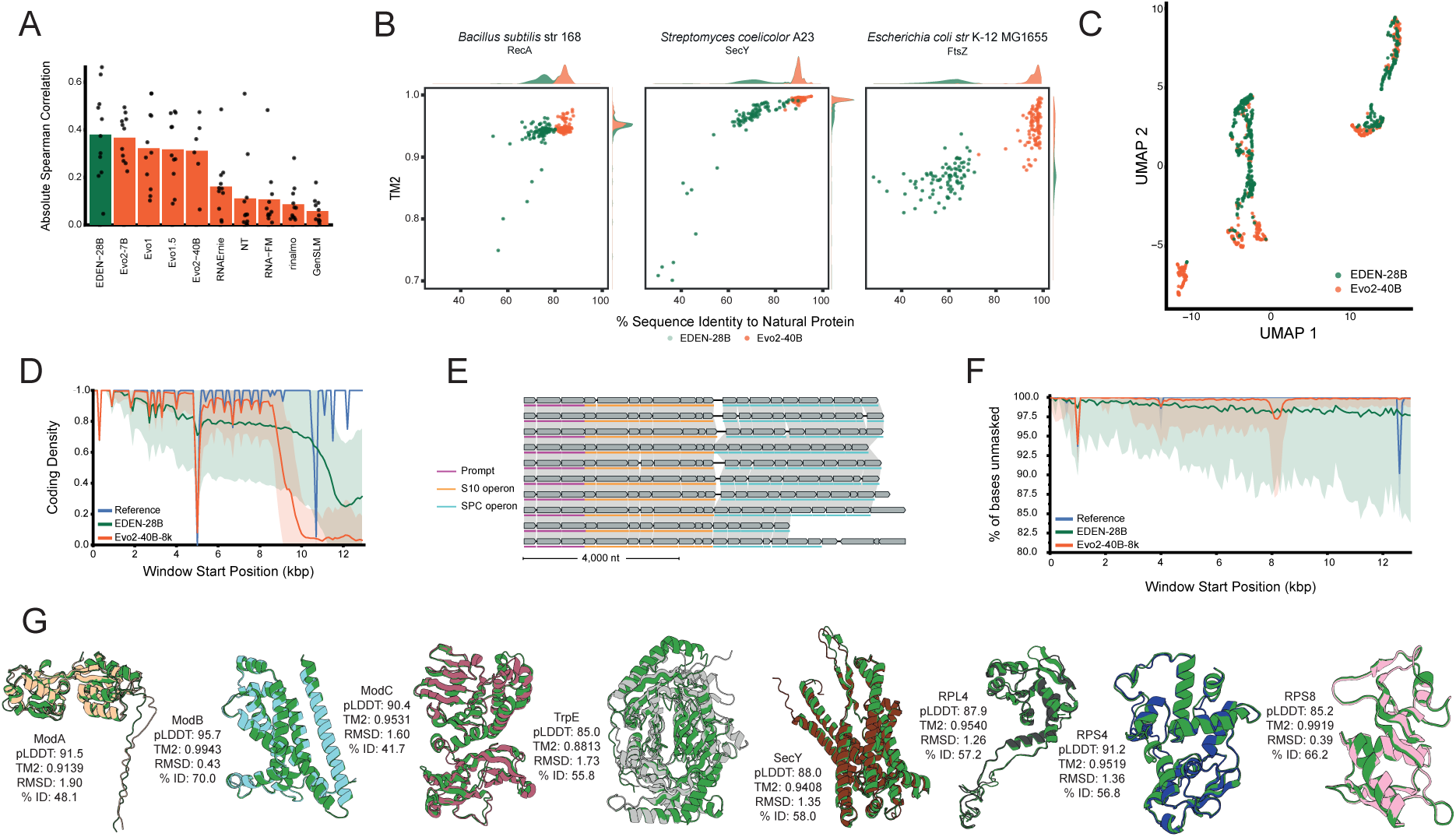
EDEN model evaluations. **A** Mutation effect prediction for prokaryotic protein coding genes based on RNAGym DMS datasets, with EDEN showing state of the art performance. **B** Protein recovery of conserved bacterial genes using a 30% sequence prompt shows high diversity in sequence space but conserved structural homology for EDEN model (n = 100). **C** UMAP of LSRs generated by EDEN and Evo2, showing both overlap and expansion across sequence space. **D** Coding density of reference genome, EDEN, and Evo2 across S10 operon covering 14,000 bp. **E** Exemplary syntenies of S10 operons and the downstream conserved SPC operon across exemplary EDEN generations. **F** Sequence complexity across 14,000 bp visualized as percentage unmasked for the reference, EDEN-generated, and Evo2-generated S10 operon (same reference and generations as in D). **G** 8 exemplary protein superimpositions between EDEN-28B generations and natural counterparts, indicating a range of sequence identities whilst maintaining high structural homology.

We moved on to evaluate EDEN’s generative capabilities, and compared these to the capabilities of Evo2 40B. First, we studied the quality of genomic generations of conserved genes from model organisms. We prompted EDEN and Evo2 40B with the 5’ end of the gene (20% of the ORF) and evaluated the ability of each model to generate the remainder. Both models demonstrated the ability to autocomplete the correct gene reliably (Figure 2B).

Notably, EDEN generated genes that were further apart in sequence space while remaining structurally consistent, with TM-score values exceeding 0.8. We also compared large serine recombinase proteins designed by EDEN and Evo2, using 30% of the 5’ end of the gene as prompts (Figure 2C), showing both overlap and unique sequence space coverage between the two generative models.

We then moved beyond single-gene generation to evaluating multiple-gene and long-context generations. Using the well-conserved ribosomal S10 operon as a case study, EDEN consistently generated sequences with coding densities greater than 0.7, beyond the 8192 nucleotide context-length used in model training (Figure 2D).

Genes generated downstream of the prompt appeared in the expected order and orientation, reflecting the canonical organization of ribosomal operons (Figure 2E). This demonstrates that EDEN can model patterns of gene synteny and operon structure from sequence data alone, without relying on explicit annotations. In addition, generated sequences maintained high DUST complexity across the entire 13kbp length (Figure 2F).

Across multiple operons tested, the model generated the subsequent proteins with high TM-scores and pLDDT values, while still retaining substantial sequence diversity (Figure 2G). Overall, the results suggest that EDEN captures information from across the dataset to generate novel diversity that preserves the structural constraints of the encoded proteins.

### AI-Programmable Gene Insertion (aiPGI) with EDEN

As discussed above, a programmable strategy for site-specific integration of multi-gene-length DNA into the human genome would enable precise replacement of pathogenic alleles with healthy copies of the gene and support the construction of complex, multicomponent genetic circuits for cell-based therapies targeting cancer and autoimmune disease.

Among existing gene-integration platforms, large serine recombinases (LSRs) are distinguished by their compact size (∼500 amino acids), the ability to insert large DNA cargos of arbitrary length in specific genomic locations without generating double-strand breaks or requiring host DNA repair pathways, a broad repertoire of target specificities shaped by extensive phage–host co-evolution, and highly reproducible integration profiles that enable systematic de-risking of off-target events^94,97^.

EDEN was pretrained on a large dataset of metagenomic sequences, including sequences that preserve a direct record of phage insertion into host genomes. These sequences provide an explicit link between phage-encoded genes required for precise genomic targeting and their corresponding DNA target sequences. In the case of LSRs, the relationship between bacterial and phage attachment sites (attB and attP, respectively) and the LSR coding sequence defines a grammar of DNA–protein interactions (Figure 3A) that, if learned, could enable an AI programmed approach to large gene insertion.

**Figure 3:**
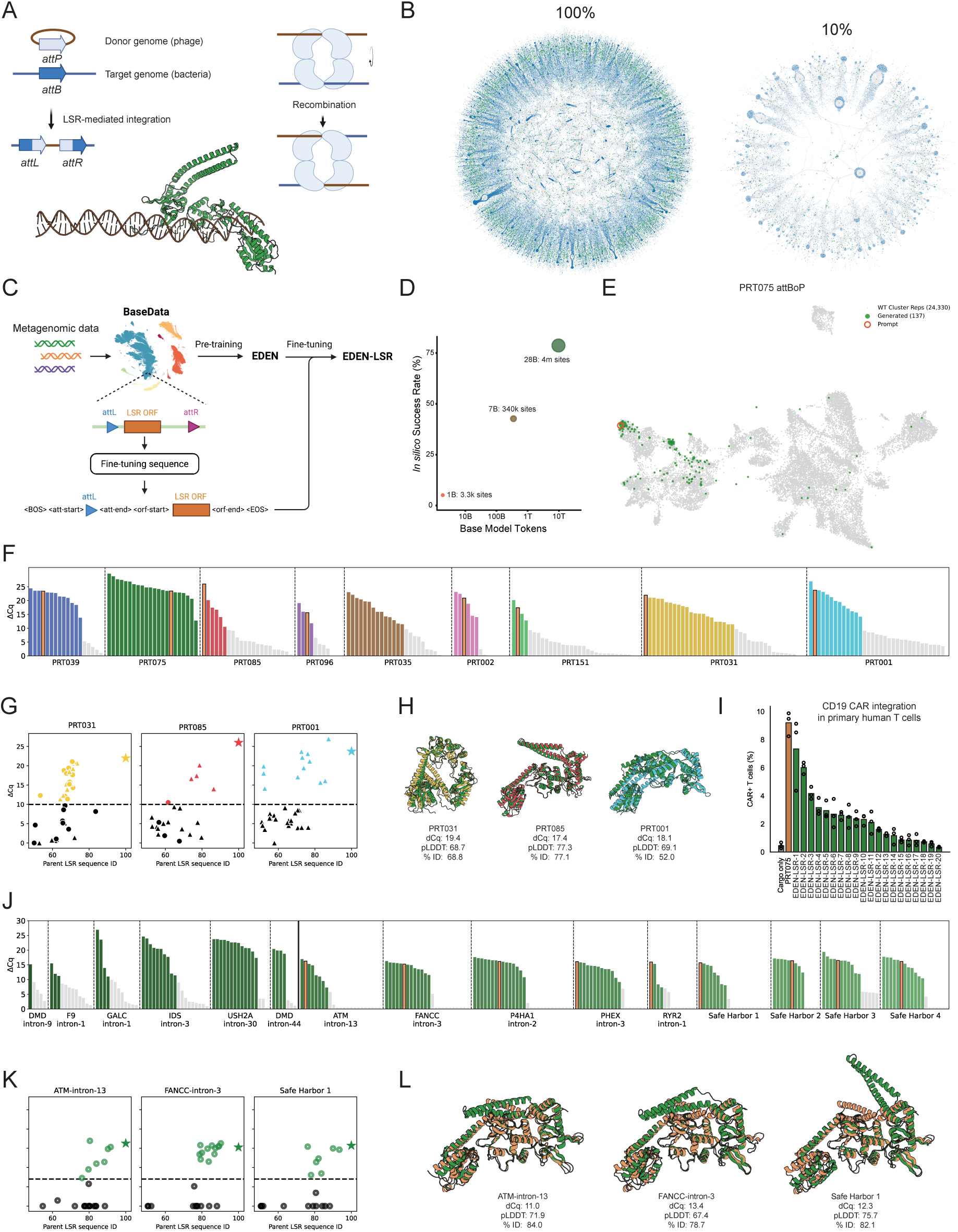
Utilization of EDEN for AI-programmable gene insertion (aiPGI). **A** Diagram of Large Serine Recombinase (LSR) mechanism including oligomerisation and a visualisation of LSR monomer bound to DNA. **B** Multi-million-node graph of LSRs (green, clustered at 80% sequence identity) connected to their paired attachment sites (blue) after bioinformatic mining. LSR dotsize (green) is proportional to the cluster size. 100% of att-LSR dataset (left) and 10% subsample of att-LSR dataset (right). **C** EDEN fine-tuning procedure yielding the EDEN-LSR model used for aiPGI applications. **D** For EDEN-LSR, scaling the basemodel token (up to 10 trillion) and parameter counts (up to 28 billion) yields higher model performance *in silico* towards better LSR generation (measured by protein domain presence) across log scales. **E** UMAP showing the distribution of generated LSRs in ESM2-embedded space when prompted on the wild-type site from one of the most abundant LSR clusters (PRT075). **F** Experimental recombination activity of 176 LSRs generated by EDEN when prompted on the wild-type sites of nine active natural LSRs (across each of the nine plots). Orange bars show wild-type LSR activity within each group. **G** Experimental activity vs sequence identity to wild-typeLSR for selected LSR prompts. AttBoP’ prompts are shown as triangles, attB-half prompts as circles, and wild-type LSRs as stars. **H** Example Boltz-folded structures of functionally active generated LSRs (various colors) superimposed with their corresponding wild-type LSRs (green), covering a range of sequence identities. **I** CD19 CAR integration in primary human T cells mediated by PRT075-based AI generated LSR variants. Integration percentage was measured by anti-CD19 flow cytometry; wild-type LSR (orange) and EDEN-generated LSRs (green). **J** Experimental recombination activity of LSRs generated by EDEN when prompted on therapeutically relevant target sites (30bp) in the human genome and not found in EDEN’s training data. First eleven plots show results from prompting on 11 loci in introns in disease-relevant genes, latter four show results from prompting on 4 putative safe-harbor sites. **K** Activity vs sequence ID to PRT075 for selected pseudosite prompts, with all LSRs generated from attB-half prompts and shown as open circles. **L** Example Boltz-folded structures of active generated LSRs (green) conditioned on pseudosite prompts superimposed with wild-typePRT075 structure (orange).

The ultimate goal of doing this is to prompt on a therapeutically relevant target sequence from the human genome and generate high performing recombinases that integrate large DNA payloads specifically at that target site.

Having first demonstrated that EDEN can generate diverse LSRs when prompted with the first 30% of a protein sequence (Figure 1H, 2C), we sought to extend this capability to the design of site-specific recombinases guided by a short DNA prompt containing only the desired genomic target site.

Although EDEN acquires broad evolutionary principles during pretraining, targeted fine-tuning enables the model to focus on specific structure–function relationships, here defined by ground-truth attachment site-LSR pairings. To this end, millions of LSR-attachment site pairs (Figure 3B) were mined from unlabelled metagenomic sequences in BaseData using a bioinformatics pipeline which yielded att sites that could further be used as reference sequences to identify additional LSR-att pairs from public databases. The resulting paired LSR-att-site dataset forms a complex sparse graph structure (Figure 3B). For fine-tuning, attachment sites were reoriented to the attL configuration (attBoP′ or attB-half) and concatenated with the corresponding LSR ORF, with control tokens inserted at the termini of both the attachment site and ORF sequences (Figure 3C).

The best-performing fine-tuned model, EDEN-LSR, was derived from the 28B-parameter EDEN architecture fine-tuned on millions of curated att-LSR sequences, until convergence on a held-out validation set of attachment site-LSR pairs. To evaluate EDEN-LSR, we assembled a benchmark comprising 46 phylogenetically distant attachment site-LSR pairs from BaseData that had been previously validated for activity in a biochemical assay. These LSRs spanned a wide range of similarity to sequences in the training set, including several with no homologs exceeding 70% sequence identity. For each benchmark example, the model was prompted with either a 60-bp core attL (attBoP′) or a 30-bp half-core attB (attB-half) sequence and tasked with generating up to 2,600 nucleotides. Generated nucleotide sequences were translated and assessed for domain content using HMMER, and all LSR-containing ORFs were evaluated for sequence similarity to both the ground-truth LSR corresponding to the attachment site prompt and the nearest homolog present in the training set.

Across model variants trained with increasing parameter counts and progressively larger pretraining and fine-tuning datasets, we observed a strong scaling relationship between model and data size and performance (Figure 3D).

Under both prompting schemes (attBoP′ or attB-half sequence), EDEN-LSR consistently generated full-length LSR ORFs, with more than 74% of all generations having the correct domain architecture (resolvase, recombinase, and zinc beta ribbon). The majority of these sequences also exhibited high predicted foldability, with ESMFold pLDDT scores comparable to those of native LSR sequences.

Next, we investigated the global conditionality of generated sequences within the natural LSR sequence space, using an orthogonal protein language model (ESM2-650M) to generate embeddings for both native and generated protein sequences and visualizing their distribution on a UMAP projection (Figure 3E). We observed that while generated sequences are frequently represented in the vicinity (>70% sequence identity) of the native LSR for the paired wild-type attachment site (5.9% of all generated sequences), many generated sequences occupy more distant regions, indicative of the broad diversity of generated LSR sequences for any given prompt. The near-native generation rate varied substantially across prompts and tended to correlate with the abundance of the corresponding LSR in the BaseData training set, as measured by cluster size (Supplementary Figure 2). For one of the most highly represented LSR clusters, here named “PRT075”, 34% of generated sequences exceeded 70% sequence identity when prompted on the wild-type attBoP′ site.

To evaluate the activity of generated LSRs under experimental conditions, we developed a rapid biochemical recombination assay, encoding LSR ORFs within double-stranded DNA fragments containing a T7 promoter and flanked by attB and attP attachment sites. Incubation of these fragments in an *in vitro* transcription–translation (IVTT) system produces the corresponding LSR protein; if active, the enzyme catalyzes recombination between the attachment sites to generate a circularized DNA product, which is subsequently quantified by qPCR.

First, we evaluated LSRs using attBoP′ and attB-half prompts derived from nine active native LSRs in BaseData. EDEN-LSR was prompted with the corresponding native attBoP′ or attB-half sequence. To prioritize EDEN-LSR-generated proteins for experimental testing, we selected *in silico* generations exhibiting 50-90% sequence identity to the native LSR associated with each prompt. We further filtered candidates to limit similarity to any sequence in the training set and to enforce diversity among generated sequences using an all-by-all distance criterion. This procedure yielded 818 candidate LSRs, from which 176 were selected for experimental testing.

In total, 53.6% of generated LSRs exhibited significant recombination activity, with similar success rates using attBoP′ and attB-half prompts (48.1% and 63.2% respectively) (Figure 3F). In this biochemical assay, across all prompts, 23 tested LSRs (13%) exhibit activity levels similar to the native LSR from the corresponding prompt. EDEN-generated LSRs also exhibit significant sequence diversity – the most divergent active LSR had only 52% sequence identity to its matched native LSR (Figure 3G). Despite low sequence identity, the generated LSRs adopt folds closely resembling the corresponding wild-type LSR structures, as predicted by Boltz-2 (Figure 3H)^52^.

To evaluate activity in a therapeutically relevant system, we selected twenty EDEN-generated LSRs based on PRT075 and evaluated them for CD19 CAR insertion into primary human T cells. Using mRNA to express the LSR and a plasmid template containing the cognate attP sequence and CD19 CAR expression cassette, we found that 50% of EDEN-generated LSRs were capable of CAR insertion into T cells (significantly higher activity than that of the empty cargo control, p<0.05), with one candidate having equivalent activity to the WT parental protein (Figure 3I). Future work is planned to investigate the specificity of these LSRs. Together, these data show that EDEN-LSR is capable of generating active, sequence diverse LSRs when prompted with wild-type bacterial att-sites, and are capable of achieving therapeutically relevant levels of gene insertion in human T cells.

Next, we set out to generate functional LSRs by conditioning the model on pseudo–attachment sites from the human genome. These pseudo-sites are not found in the training data and include sites in disease-relevant genomic locations. We focused on a native LSR from BaseData (PRT075) that exhibited a high success rate with the native attB prompt (Figure 3F), and demonstrated therapeutically relevant levels (∼40%) of integration in human cells (K562) (Supplementary Figure 3). We identified fourteen therapeutically relevant target pseudosites for this LSR spanning a range of genomic contexts, including 4 putative safe-harbor and 10 loci in early introns in disease-relevant genes (ATM, DMD, F9, FANCC, GALC, IDS, P4HA1, PHEX, RYR2, USH2A). These pseudosite sequences had a 54-75% sequence identity relative to the closest native bacterial attB site in the training data. We prompted EDEN-LSR with the 30-bp attB-half sequence of these therapeutically relevant pseudosites and sampled approximately 10 generated sequences per target for experimental testing, using a sequence identity threshold of >50% vs native PRT075 to select candidates. Pseudosites were then tested for recombination with the native attP sequence.

Notably, despite being prompted using only a short (30bp) DNA target sequence completely outside the training data, EDEN generated multiple successful LSRs for every targeted pseudosite (Figure 3J). For the most successful prompts (safe harbor 2-chr13, safe harbor 3-chr13, and safe harbor 4-chr8), 66% of tested LSRs had significant recombination activity on the pseudosite; others exhibited a lower success rate, with 20% of LSRs generated from DMD-intron 9 displaying significant activity. Across all prompts, 48% of the generated LSRs tested positive for recombination on the corresponding pseudosite, with 27 LSRs (16%) exhibiting activity similar to the native PRT075, including proteins with as low as 76% sequence identity to PRT075 (Figure 3K). Further work is planned to validate the activity of these psuedo-site prompted LSRs in relevant human cell models, building on the success in cells of the wild-type prompted LSRs above.

In summary, these results demonstrate that EDEN-LSR is capable of generating diverse, active LSRs when prompted directly with short (30bp) genomic target sites, including therapeutically relevant sequences from the human genome that were absent from the training data. As natural LSRs in BaseData have been experimentally shown to integrate at over 10,000 disease-relevant sites in the human genome (data not shown), this establishes a powerful new approach for engineering therapeutic LSRs with activity at defined genomic sites.

While initial findings indicate successful on-target activity, realizing the full potential of safe, programmable recombinases for large-payload medical applications will require further model optimization and experimental validation. Future work will integrate reinforcement learning to refine control over activity and specificity, alongside comprehensive assessment of integration efficiencies and off-target profiles in relevant human cell populations.

### EDEN designs active and novel bridge recombinases

In contrast to LSRs, where DNA targeting is programmed entirely by the amino acid sequence of the protein, bridge recombinases (BRs) offer an RNA-programmable protein complex that is capable of genetic insertion and excision in the context of the human genome^129^, positioning it as an emerging gene editing technology for potential therapeutic applications.

The BR system consists of a transposase protein of diverse classes (of which the IS110 and IS1111 classes are the most well-studied), and a ncRNA element, termed bridge recombinase guide (bRNA), that is either upstream (i.e. left element or LE) or downstream (i.e. right element or RE) of the transposase (Figure 4A). The bRNA folds into a secondary double-loop structure, each of which binds to the donor and target sequence based on sequence-specific guide motifs^130,131^. Physical interactions between the BR and the bRNA ultimately facilitate the joining of the target and donor sequences and subsequent recombination events^132^.

**Figure 4:**
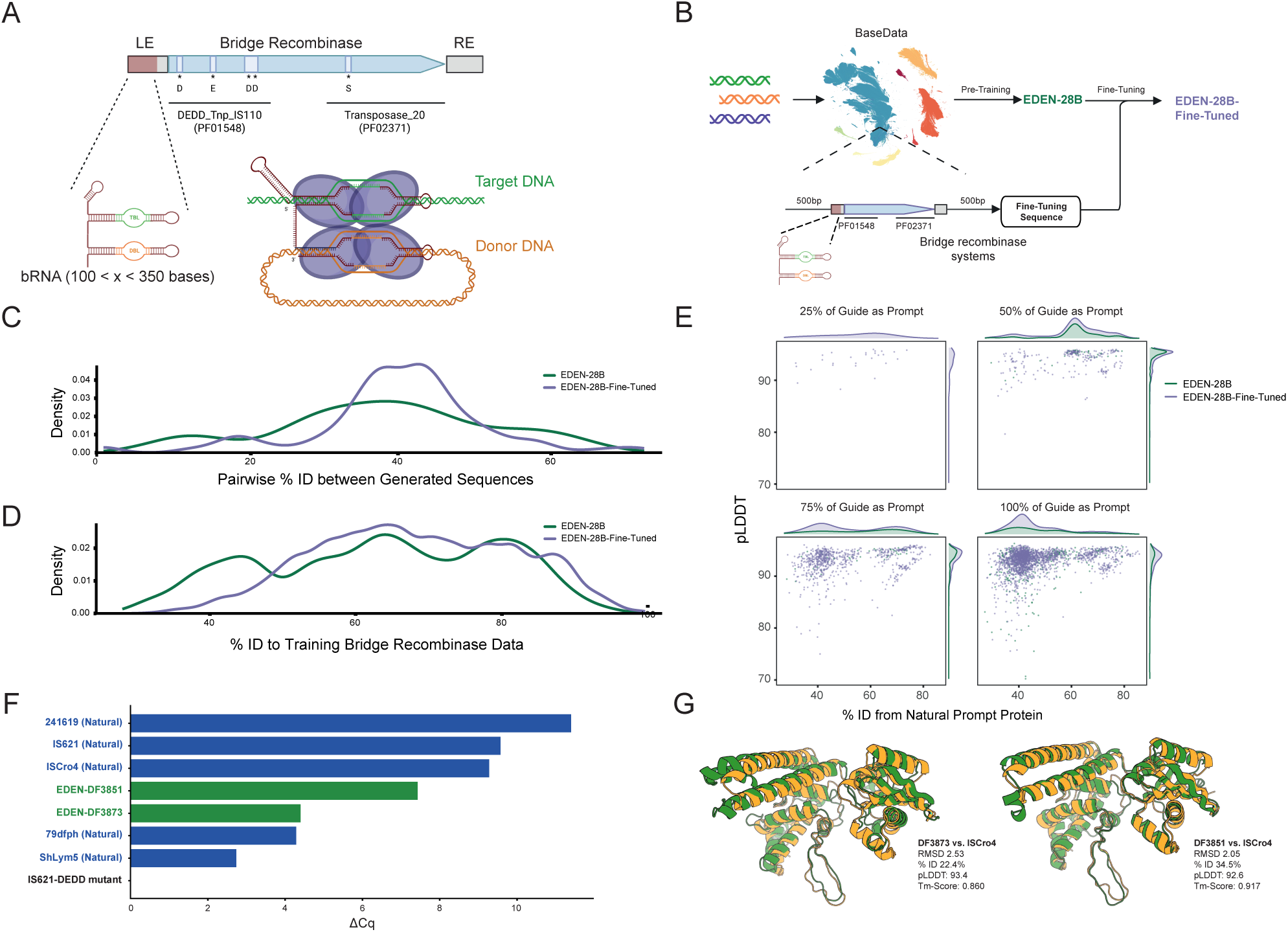
EDEN designs active and novel bridge recombinases. **A** Diagram of bridge recombinase system displaying the bridge RNA, target DNA, donor DNA, and recombinase components. **B** Fine-tuning strategy using EDEN-28B and BaseData bridge recombinase systems for EDEN-BR. **C** Density of pairwise sequence identity of EDEN-generated bridge recombinases. **D** Density of sequence identities of EDEN-generated bridge recombinases compared to training data. **E** pLDDT and sequence identity distributions of EDEN-generated BRs for different fractions of guide RNA prompts. **F** IVTT assay results of EDEN-generated and wild-type BRs. **G** Structural superimpositions of EDEN-generated, active BRs compared to ISCro4.

The EDEN-28B base model demonstrates that prompting with as short as 50% of the upstream guide RNA sequence was sufficient to generate the downstream recombinase gene encoding BR proteins unique to public and BaseData natural sequences. These generated proteins bear the RuvC-like domain with the DEDD catalytic residues, as well as the Tnp domain with its conserved serine residue.

EDEN-BR was created by fine-tuning the base model on over six million BR-containing genomic regions from BaseData, (Figure 4B and Methods). EDEN-BR improved the generation of BR proteins by over eight-fold when the complete guide was used as prompt (Supplementary Figure 4).

For 99 bridges the fine-tuning increased the number of guide prompts giving rise to downstream coding sequences with the expected functional annotations. At the same time, within a given prompt, fine-tuned generations maintained a spread of diversity with respect to other generated BR proteins (Figure 4C). Significantly, the fine-tuned EDEN-BR model generated more diverse BR proteins with reference to those observed in the training data (Figure 4D), as well as the natural recombinase protein associated with the bRNA prompt that was used in the generation (Figure 4E). Overall, for both base and fine-tuned models, generated BR proteins maintained highly confident structure predictions despite the wide sequence diversity spread (Figure 4E). Pilot biochemical, cell free validation assays were conducted on a small set of 49 sequences generated by EDEN-BR with 100% length bRNA. These tests demonstrated recombinase activity for two sequences, named DF3873 and DF3851 (Figure 4F).

These two active AI designed BRs are no more than 85% identical in sequence to any BaseData or public sequences (Table 1). Importantly, these generated proteins present high structural homologies to ISCro4, the best characterized BR protein to date^129^, despite being no more than 35% similar in sequence to ISCro4 (Figure 4G).

**Table 1.** Sequence identity between DF3873/DF3851 and BaseData and public databases.

| Database | Closest Match to DF3873 (%)* | Closest Match to DF3851 (%)* |
| --- | --- | --- |
| **BaseData** | Proprietary Sequence A (84.7) | Proprietary Sequence B (82.4) |
| **NCBI** | HMK23979.1 (57.9) | MBW2541154.1 (55.0) |
| **MGnify** | MGYP003941236639 (57.6) | MGYP003449954008 (79.9) |
| **UniProt** | A0A2U3KKS4 (57.8) | A0A2Z5GAY9 (52.2) |
| **EMBL Patent** | USPTO:WFL37734 (40.2) | USPTO:AAO95525 (33.6) |

\*hits above 90% coverage based on query sequence

Overall, we show successful design of novel, active bridge recombinases by EDEN when only prompted on the non-coding bridge RNA sequence, and in some cases, even only a fraction thereof. We further show significant divergence of the EDEN-generated BRs from both training data as well as public BRs. Viewing these as an orthogonal approach to LSR design for programmable gene insertion, we show that EDEN is a foundation model capable of designing candidate therapeutic molecules across different modalities and protein families.

### EDEN designs potent antimicrobial peptides

First, we screened BaseData for antimicrobial peptide (AMP) activity to confirm that the pretraining data contained active AMPs. For this, we conducted a search for small open reading frames (smORFs) with all assembled sequences in BaseData. After size filtering, over 300 million peptide sequences were evaluated using the APEX pathogen prediction model^133^, which generated minimum inhibitory concentration (MIC, µmol L⁻¹) predictions across a training set of pathogens; lower MIC values correspond to stronger inhibitory activity. The median MIC across individual pathogen-specific predictions was used for downstream analysis (Figure 5B). Using a cutoff of 64 µmol L-1, the dataset was prioritized to over 20,000 sequences. These candidates were further analyzed through comparisons to the DRAMP v4.0 database^134^, taxonomic annotation, physicochemical property profiling, and representative clustering. One peptide could not be synthesized; however, all remaining 34 candidates inhibited growth across a panel of 20 gram-positive and gram-negative pathogenic strains, with MIC values ≤ 64 µmol L⁻¹. Notably, several candidates exhibited strong activity at low concentrations (<2µmol L⁻¹) against approximately 80% of the tested strains. (Supplementary Figure 5).

**Figure 5:**
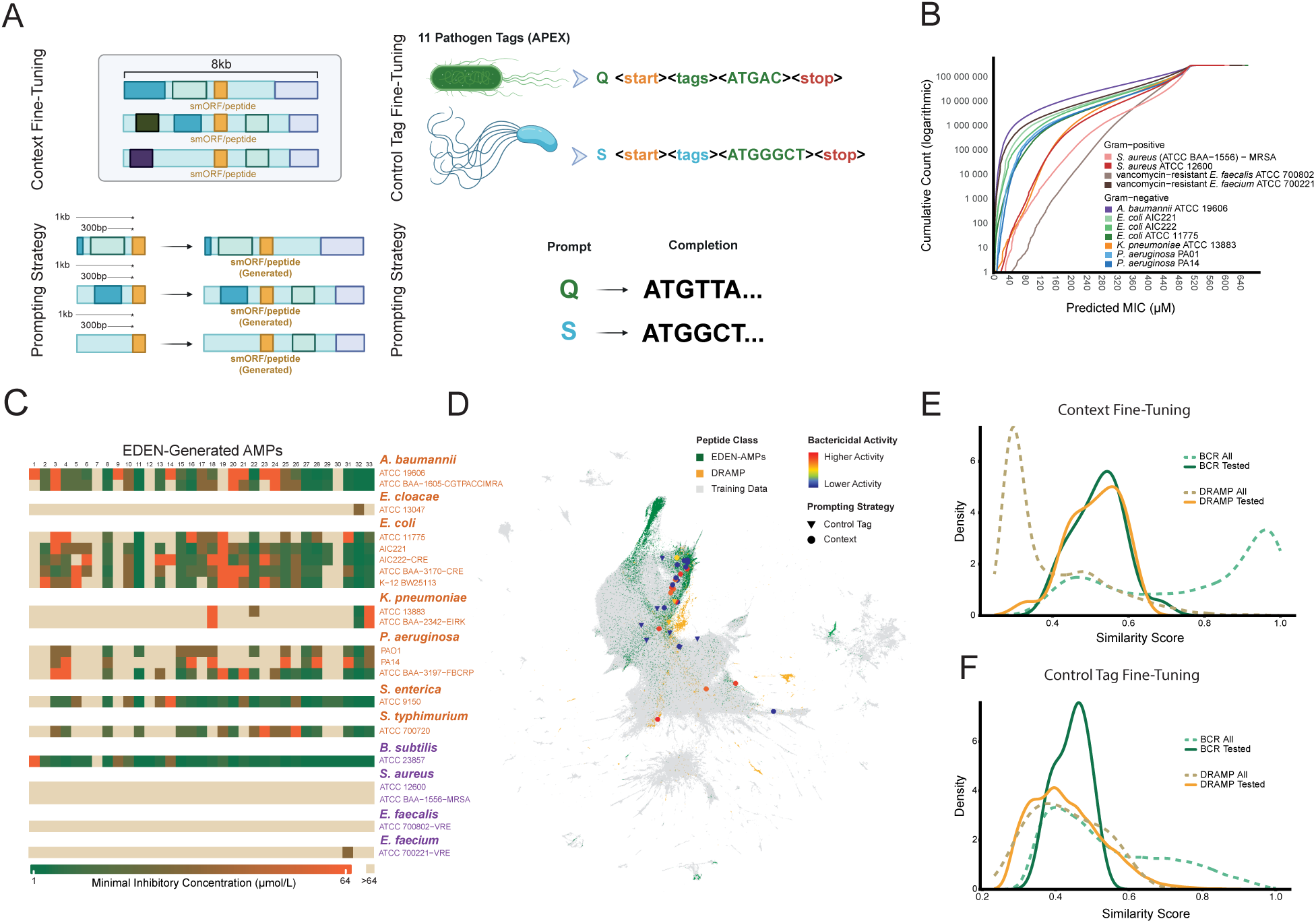
EDEN designs potent antimicrobial peptides. **A** Fine-tuning and prompting strategy for antimicrobial peptide generation. **B** Predicted individual pathogen MIC values across all smORFs in BaseData using APEX pathogen. **C** Heatmap showing the results of activity validation assays confirming antimicrobial activity of EDEN generated peptides against 16 clinically relevant pathogen strains. **D** UMAP visualization of EDEN generated AMPs against the training data and DRAMP. Activity range is the same as indicated in subfigure C. **EF** Sequence similarity distributions of generated peptides relative to BaseData and DRAMP. Context fine-tuning yields higher similarity to reference datasets, whereas control-tag fine-tuning produces more divergent, novel sequences.

We then developed a fine-tuning and prompting strategy in order to generate synthetic AMPs (Figure 5A). Leveraging EDEN-28B, we developed two fine-tuned models using distinct but complementary datasets, built from analyses and results produced from the natural BaseData AMP activity exercise. The first, a context fine-tuned model, was trained on genomic sequences capturing the local contextual neighborhoods of smORFs encoding putative AMPs. This training set comprised over 15,000 sequences, totaling approximately 115 million tokens.

For the second model, we introduced control tags. This was trained on peptide sequences represented with explicit start and stop tokens and prefixed pathogen-specific control tags to enable conditional generation. Control tag conditioning has been explored in generative language modeling and is emerging in biological sequence models to steer generation toward desired properties^135^. The control-tag training dataset comprised over 1.5 million sequences, totaling approximately 64 million tokens.

Prompting strategies for both fine-tuned models are shown in Figure 5A. For the context fine-tuned model, prompts of varying lengths derived from sequence upstream of the smORF were used to generate over 80,000 peptide sequences. The control tag fine-tuned model generated over 400,000 sequences by prompting on pathogen-specific control tags or in an unconditional manner using the start tag alone.

Given the scale and diversity of the generated sequences, additional *in silico* selection was required prior to experimental testing. We therefore subjected the predicted AMPs to further confirmation analyses. Leveraging predicted MIC scores on the generated sequences using APEX pathogen, we first showed that effective antimicrobial peptide design can be achieved with EDEN directly from a genomic context. Additionally, by adding predicted pathogen tags to the training data, we generated peptides with lower predicted MIC values for certain pathogens including increasing the number of predicted AMPs by approximately 16-fold for *E. coli* ATCC 11775 compared to unconditional generation. We were able to generate predicted AMPs for 10 out of 11 strains. In particular, generation of AMPs predicted to be activated against *A. baumannii* ATCC 19606 was highly successful, with over 15,000 predicted AMPs generated across both fine-tuning strategies. Through the two models we have built a collection of predicted AMPs of over 25,000 sequences using a stringent activity threshold of predicted MIC of ≤32 for at least one pathogen and a length threshold of 8 - 50 amino acids.

To assess novelty, we compared the predicted AMPs to known AMP sequences using sequence similarity metrics. By calculating a Striped Smith-Waterman similarity score to the natural AMPs for the fine-tuning dataset of each model and the DRAMP database, we were able to confirm substantial novelty to both the training data and publicly available AMPs (Figure 5E, 4F). In particular, a large proportion of EDEN generated peptides have a similarity of below 0.7 to any natural AMP. These results indicate that the generated peptides are not close variants of sequences in the training data or public AMP databases, but instead exhibit low pairwise similarity to known AMPs.

Additionally, the EDEN generated predicted AMPs exhibit amino-acid compositions and physicochemical property distributions that globally overlap with those of natural AMPs (Supplementary Figure 6A-E). In particular, a substantial fraction of peptides fall within the canonical cationic charge range associated with AMP activity (+2 to +9). For example, 49.6% of peptides in the DRAMP reference set and 58.7% of fine-tuning peptides fall within this range, compared with 28.0% of the EDEN generated predicted AMPs. While the charge distribution of generated peptides is not identical to that of natural AMPs, this partial overlap indicates that EDEN captures key physicochemical trends relevant to antimicrobial function. This is consistent with a noticeable enrichment of lysine (K) in comparison to natural AMPs, a pattern previously seen in generated peptides^136^.The hydrophobicity distributions of the generated peptides overlap substantially with those of natural antimicrobial peptides. DRAMP peptides show near-neutral median hydrophobicity (median −0.02), while fine-tuning peptides (median −0.19) and Eden-generated peptides (median −0.41) span a similar overall range of values. Although the generated peptides are modestly shifted toward lower hydrophobicity, their distributions retain broad overlap with natural AMPs.

ESM650M embeddings were used to project the predicted AMPs into UMAP space alongside AMPs from the fine-tuning dataset and DRAMP (Figure 5D). The generated peptides broadly overlap with known antimicrobial sequence space while also extending into less densely populated regions, indicating that EDEN can produce sequences consistent with established AMP characteristics while introducing sequence diversity. EDEN therefore recapitulates key chemical features of natural AMP distributions while significantly expanding coverage and sequence novelty beyond known examples from public data. Both the similarity scores and the UMAP projections demonstrate that the sequences exhibited substantial novelty, highlighting the model’s ability to design towards broader sequence space and enable realistic yet diverse AMP generation.

To test the potency of EDEN’s AMP generations we synthesized and tested 33 generated AMPs *in vitro* against a panel of pathogenic bacteria. The panel consisted of six gram-negative species (*Acinetobacter baumannii*, *Escherichia coli*, *Klebsiella pneumoniae*, *Pseudomonas aeruginosa*, *Salmonella enterica*, *Enterobacter cloacae*) and four gram-positive species (*Staphylococcus aureus*, *Bacillus subtilis*, *Enterococcus faecalis*, *Enterococcus faecium*), and included both drug-susceptible strains and multidrug-resistant clinical isolates.

97% of the selected EDEN-generated peptides demonstrated antimicrobial activity inhibiting bacterial growth at concentrations ≤64 µmol L^-1^ in wet-lab assays, confirming that EDEN’s *in silico* designs translate into experimentally validated function (Figure 5C). In particular, several generated AMPs showed broad activity across multiple strains, with 27 AMPs active across five or more strains. The antimicrobial activity of the tested AMPs was stronger across the gram-negative strains, and all *Acinetobacter, Escherichia, Klebsiella, and Salmonella* strains had MICs of 1 - 4 µmol L^-1^ for at least one tested AMP. This observation is of particular interest, as gram-negative bacteria present unique structural barriers to AMPs, including a lipopolysaccharide-rich outer membrane and specialized resistance mechanisms that impede peptide permeation and activity, rendering them intrinsically more difficult targets than gram-positive organisms^137^. The tested peptides covered diverse regions of the AMP sequence space and were highly divergent, all tested sequences had a similarity score below 0.7 and strong activity observed for sequences exhibiting high novelty (similarity < 0.4).

In summary, we show that EDEN can generate potent antibiotics for a range of drug-resistant pathogens, significantly expanding beyond the training data and publicly known peptides. To our knowledge, this marks the first instance a DNA foundation model has been used directly for peptide and antibiotics design with proven potency in ground-truth experiments against targets of interest.

### EDEN designs multi-species synthetic microbiomes

With previous generative biological foundation models showing successful design on the open-reading frame, mobile genetic element, and genome level^7,123^, we sought to leverage EDEN for design tasks beyond individual genomes and generate a fully synthetic microbiome. The motivation behind this lies in the recent development of therapeutic solutions in chronic diseases through the engineering of microbial consortia for chronic diseases such as immune-mediated colitis^138^. To assess this design potential, samples in BaseData were characterized to find annotation signatures specific to the biome where it was collected. This was conducted to curate a set of biome specific prompts, create a baseline of metrics used to evaluate sequences and to identify a well characterized sample for fine-tuning (Figure 6A-B).

**Figure 6.**
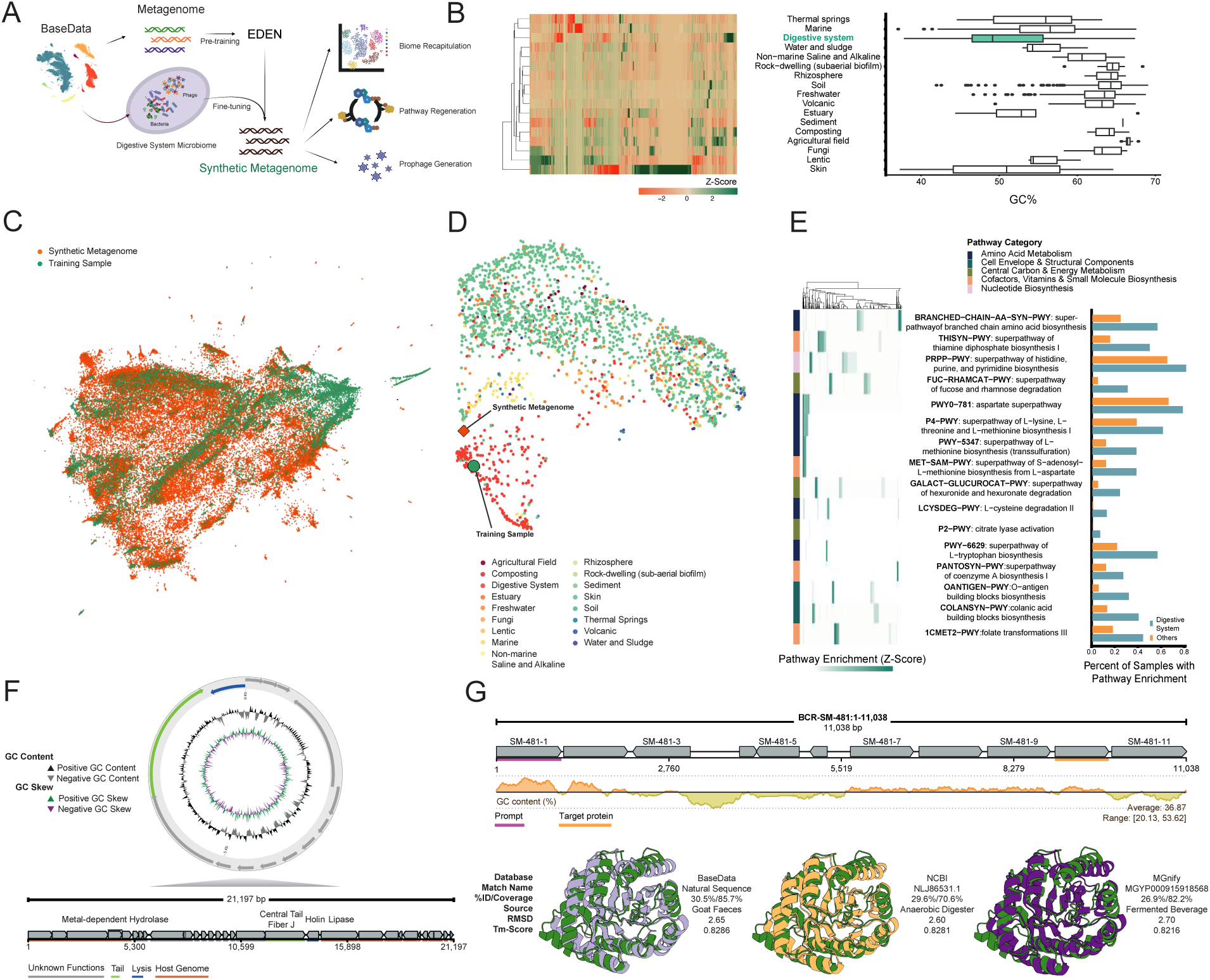
Towards multi-species synthetic microbiomes. (A) Strategy for generating a synthetic metagenome. BaseData™ was used to identify appropriate sample(s) of digestive systems which was used to fine-tune the EDEN base model. This fine-tuned model was used to generate a population of sequences seen in a digestive system sample. (B) All samples from biomes in BaseData™ were characterized to identify KEGG annotation signatures for use in downstream evaluation metrics and to construct appropriate prompts. Shown are biome wise scaled KEGG gene enrichment and GC%. (C) UMAP projection of training and synthetic metagenome sequences based on k-mer composition, illustrating their distribution and overlap in reduced dimensional space. (D) UMAP of BaseData™, training sample and synthetic metagenome sample based on Jaccard distances of taxon presence–absence profiles, colored by biomes. (E) Overview of a selection of top 16 metabolic pathways significantly enriched in the synthetic metagenome, showing scaled counts of pathway genes between sequences (left) and their corresponding natural enrichment in BaseData™: digestive systems versus other biomes (right). All 16 pathways show ≥ 60% completion in the generated synthetic metagenome (F) A generated sequence from the synthetic metagenome outlining a prophage-like sequence and its genetic architecture. (G) An 11 kb generated genomic sequence encoding multiple ORFs, highlighting the tenth ORF with minimal detectable sequence homology to the training set and public databases. Despite low sequence identity, the predicted protein adopts a well-formed fold with high structural similarity, indicating preservation of genomic context beyond the prompt region.

The specific biome that was chosen for this assessment was “digestive system” (following MGnify biome hierarchy) due to its unique set of KEGG annotations and well defined sequence profile (Figure 6B)^82,139^. Digestive systems were particularly enriched in KEGG annotations such as K04029 (ethanolamine utilization protein) and K02791 (maltose/glucose PTS system EIICB component), had a GC profile range between 47-55% (Figure 6B), and a coding density of approximately 82%, which is a the slightly lower end of the traditionally reported prokaryotic coding density range of 80-95%^140,141^. A single digestive system microbiome sample containing over 160,000 contigs was selected from BaseData and used to fine-tune EDEN for 1.5 epochs. We designed prompts based on the context that is adjacent or at digestive system annotations specifically extracting sequences of length ranging 500 bp to the length of a full gene. Using a vLLM inference engine (Methods) paired with the fine-tuned model, these prompts generated more than 100,000 sequences of 10 kb each, totaling a gigabase scale synthetic microbiome (Figure 6A)^142^.

We then characterized the synthetic microbiome at a global scale according to sequence space and taxonomic constitution. Encouragingly, the synthetic microbiome generated by EDEN recapitulates and expands beyond the sequence space of the digestive system fine-tuning sample (Figure 6C). When annotating the synthetic microbiome with kraken2, we identified 9067 taxonomic units, 7533 (83%) of which were shared with the sample used for fine-tuning and 9006 (99%) were shared with other samples from the same biome, suggesting that EDEN, when generating the synthetic microbiome, captures cross-species biological consistencies beyond the explicit taxonomic make-up of the fine-tuning data. When representing the synthetic sample within a UMAP with samples from BaseData based on taxonomic breakdown, the synthetic microbiome is placed in taxonomic space consistent with natural digestive system samples (Figure 6D).

In order to study *inter*-and *intra*-genomic functional features of the synthetic microbiome, we analysed metabolic pathway abundance across natural samples, identifying a set of 16 pathways with significantly higher proportions in digestive system samples compared to others (Figure 6E). We further show that for all of these 16 pathways identified, we observe a significant enrichment of these within the synthetic microbiome, all of which at completion rates above 60% (Figure 6 E). We note that several of the pathways listed here are commonly completed across, and not within species in microbiome environments, such as the superpathway of coenzyme A biosynthesis^143^.

A metagenomic sample represents a population of organisms at a given time. This may include but is not limited to microeukaryotes (in environmental samples), bacteria and bacteriophages. We observe that a portion (21,561) of our generated and filtered sequences have been annotated as phage using geNomad^144^, including sequences that may contain phage fragments. We also observe prophage annotations, where geNomad has annotated a portion of the sequence as a prophage, and the remaining sequence as host. We performed a second round of generations (Methods) producing a prophage annotation flanked by its host genome, indicating that the underlying phage architecture is preserved rather than fragmented (Figure 6F). This is confirmed using geNomad annotating a prophage between the region coordinates of 5 kbp and 12 kbp within the 21 kbp generated sequence with a score of 0.9988. It also classifies the prophage sequence as *Caudioviricetes*, a common class of phages found in gut metagenomes^145^.

Whilst our prophage annotation has a confident geNomad score, it is important to note that geNomad, like many common prophage annotation tools, approximates boundaries of prophages but does not identify exact start and end attachment sites^144^.Thus we also investigated the prophage genome architecture. Gene prediction tools such as Prodigal, Glimmer and GeneMark failed to correctly predict any genes in a synthetic phage genome^123^, so we used pharokka^146^ and phold^147^ to annotate genes and determine more precise prophage-host boundaries. We identify phage tail and holin gene on the prophage (Figure 6F). Holins are small hydrophobic proteins common in bacteriophages, especially tailed double-stranded DNA phages^148^ like those found in the *Caudioviricetes* class. These tools did not annotate all ORFs and a diamond blast against the NCBI database revealed low identity, high coverage matches to proteins found in gut metagenome samples. To confirm whether the host sequence flanking the prophage came from the same organism, we ran Kraken2 on the sequence upstream and downstream the prophage and they were both classified within the *Streptomyces* genus. We foresee further analysis of a *de novo* generated phage like this would benefit from likely even more sensitive functional annotation methods to have all required phage ORFs fully annotated.

In addition to demonstrating that EDEN generates functional proteins and mobile genetic elements, we further assess whether the model can preserve a biome-aware biological signal across longer context when prompting with only a sequence fragment. Using the same prompting strategy as indicated above, gene prediction/ORF calling was run on the generated sequences. After filtering by length, sequence complexity and presence of metabolic pathway annotation, a subset of the ORFs were mapped against BaseData, NCBI protein, UniProt and MGnify databases to evaluate the closest member using strict identity and coverage thresholds^60,82,149^. To illustrate candidates with low sequence homology but functional and biome-related conservation, we investigate further BCR-SM-481-10, an ORF of length 286 aa. BCR-SM-481-10 was called at 8938 to 9798 of an 11.3 kb generated sequence, representing the 10th ORF out of 11 predicted genes (Figure 6G).

To understand whether BCR-SM-481-10 was generated with a level of variability not found in the training dataset, an alignment against BaseData revealed the closest match to a protein sourced from an animal faeces sample with sequence identity of 30.5%. We characterize BCR-SM-481-10 on the sequence level revealing a xylose isomerase-like TIM barrel PFAM domain but no KEGG annotations. TIM barrels (triose-phosphate isomerase) are one of the most common structural motifs found in enzymes^150^. We used ESMFold to predict the structure of BCR-SM-481-10 (Figure 6G) to reveal a structure highly similar to the canonical TIM barrel like structure with 8 sheets in the barrel^66^. We then compared BCR-SM-481-10 with public databases. The top match from any public database was a protein from a *Firmicutes* bacterium taken from an anaerobic digester metagenome sample (ncbi) with a sequence identity of 29.6%. The fact that the top matches originate from anaerobic, digestive system environments supports the idea that the model is capturing contextual signals, consistent with BCR-SM-481-10 being generated from a digestive system derived prompt (Figure 6G). Taking this top match, we structurally aligned the ncbi hit to BCR-SM-481-10 using TMAlign and visualized using Pymol. Both matches showed strong alignment with TM scores at 0.83 (Figure 6G).

Overall, we show that EDEN, when generating long context sequences from a digestive-system prompt, produces ORFs with low sequence identity to public databases yet with structurally plausible folds, and importantly, maintains beyond-species, biological contextual signals far beyond the prompt region. Given the large number of sequences and variety of analytical approaches that can be applied to a system like this, the EDEN-generated synthetic microbiome will be further analyzed and characterized with additional taxonomic (such as marker gene based annotation^151,152^), functional, and genomic insights. The concept presented here could point towards a proof-of-principle deserving of a) further validation, including that of experimental nature, and b) broader applicability towards different biomes.

## Discussion

In this paper, we set out to evaluate the hypothesis that progress towards true programmable biology will require expanding the training datasets of generative models to include increasingly large quantities of diverse evolutionary data, far beyond the constraints of current publicly available resources. If this hypothesis is true, we would expect these models to learn increasingly universal design principles from this data and progressively improve the predictability, accuracy, and controllability of the computational design of biological code.

To test this hypothesis, we build on our previous publication of BaseData^1^ and introduce the EDEN (Environmentally-Derived Evolutionary Network) family of foundation models, the largest of which was trained on 9.7 trillion evolutionary nucleotide tokens from BaseData^1^, with no human, lab or clinical data in the pre-training dataset. In this paper, we demonstrate EDEN’s capacity for programmable therapeutic design by challenging a single architecture to design biological novelty across three distinct therapeutic modalities, disease areas and biological scales: (i) large gene insertion, (ii) antibiotic peptide design, and (iii) microbiome design.

As a result of this, we observe three key points worthy of a more extensive discussion:

(1) the EDEN training data and how training was conducted; (2) the various therapeutic applications, their successes and their limitations; and (3) the wider implications of these findings for the field.

### The role of evolutionary data in biological foundation models

Our results suggest that the path towards programmable biology lies in altering both the nature and scale of the pre-training and fine-tuning data of generative models, combined with scaling of the models themselves.

The EDEN model family has been developed in a wider field of recent and impressive achievements, but a gap remains in the training of models on data representing cross-species selective & evolutionary pressures that give rise to therapeutically relevant molecule classes at the 10 trillion token-scale and beyond. For instance, Evo2 introduced a 40B-parameter model with a 1 million basepair context window that achieves remarkable long-range genomic modeling capabilities across prokaryotes and eukaryotes^7^. Evo2’s 9.3 trillion token dataset contained ∼854 billion tokens (9%) from metagenomic sources, with the vast majority derived from eukaryotic reference genomes (e.g. >5 trillion tokens from *Animalia* alone). Similarly, while the gLM2 model was explicitly designed for metagenomics using the 3.1 trillion basepair OMG corpus, the resulting 650M parameter model was trained on ∼315 billion tokens - a training corpus similar in scale to the metagenomic portion of OG2^81^.

EDEN complements this work with a deliberate and exclusive focus on large-scale novel evolutionary data from metagenomic sources-training on 9.7 trillion tokens derived entirely from BaseData. By incorporating environmental and host-associated DNA, phage sequences, mobile genetic elements and transposons often absent from curated reference genomes at the trillion-token scale, we imbued EDEN with an especially rich repertoire of evolutionary mechanisms (such as phage-host interactions and microbiome-derived antibiotic genes) that other biological training datasets may capture to a lesser extent.

This focus on evolutionary diversity capturing inter-species signals also yielded an important insight regarding quality-aware scaling laws for biological foundation models. We observed that models trained on BaseData exhibited a steeper scaling exponent compared to those trained on curated public metagenomic datasets, achieving lower perplexity as compute increased. We attribute this to the higher quality of the evolutionary signal in BaseData - specifically the preservation of longer-range inter and intra genomic context. This provides validation to our hypothesis that expanding training data to include high-quality, diverse evolutionary context allows models to extract more useful information per token, thereby improving the predictability of biological design.

Since the EDEN models are trained on BaseData, we note that they have several key features that can be considered unique or rare in biological foundation model development: orthogonality, consistency in data collection, and consent & data governance. First, as previously stated, BaseData contains over a million previously undiscovered species, demonstrating that state-of-the art biological foundation models can be built on the basis of novel and orthogonal data outside of what is publicly known about the tree of life^1,87^. Second, each nucleotide token used to train EDEN has been derived from rigorously consistent data collection, sequencing and bioinformatics annotation pipelines which contrasts with the vast range of methodologies and protocols used in the compilation of public sequence databases^80–82,149^. Third, and uniquely for a model of this scale, each token used for pretraining EDEN has been derived from explicit stakeholder consent and benefit sharing agreements, establishing a new standard for ethical AI development in biology (see Stakeholder Best Practices section below).

Finally, while we acknowledge the utility of megabase-scale contexts for whole-genome modeling, we found that a context length of 8,192 tokens was sufficient to capture the necessary functional logic for our applications. By evaluating different options for the context length during pretraining we note the quality of generations remained high for several thousands of basepairs beyond the context length of 8,192 tokens (Figure 2). EDEN consistently maintained correct gene architectures and operon synteny (not just sequence quality or ORF density) beyond 10,000 basepairs and successfully generated complex systems like prophages by using selected regions of previous generations as prompts. This suggests that for programmable therapeutics, where the goal is often the precise engineering of functional modules in the context of accurate biological grammar, rather than entire chromosomes; a targeted, high-quality, multi-gene context window offers an efficient path to predictable biological novelty.

### Programmable therapeutic design across modalities, biological scales and disease areas

To evaluate the therapeutic utility of training models on this expanded scale of evolutionary data distribution, we tested EDEN on a series of therapeutically relevant design tasks (large gene insertion, antibiotic peptide design, and microbiome design) and demonstrated a range of new capabilities with major potential medical implications. In the context of AI-based, generative approaches for gene editing, previous work has shown impressive design outcomes, for example generating active Cas9 nucleases^6,77^. We complement these successes by using EDEN to design large serine recombinases (LSRs) and bridge recombinases (BRs) to enable programmable large gene insertion. We show that EDEN solves the inverse design problem outlined earlier in this paper by mapping directly from DNA target to a functional protein: successful designs only require prompting with the desired attachment site (in the case of LSRs) or non-coding bridge RNA (in the case of BRs).

We show that, when prompted with only 30 nucleotides of DNA representing the desired human attB genomic target site, EDEN generates multiple active LSR proteins for all tested disease-associated human genomic *loci* (ATM, DMD, F9, FANCC, GALC, IDS, P4HA1, PHEX, RYR2, USH2A) and four potential safe harbor sites in the human genome.

We further demonstrated the EDEN models’ flexibility by designing active Bridge Recombinases using only non-coding guide RNA sequences as prompts. These generated enzymes exhibited sequence identities as low as 65% relative to training data, confirming that EDEN generates novel functional machinery rather than merely retrieving memorized sequences.

Prior research and reviews have discussed the potential for AI to provide a solution for the repurposing of drugs across several rare diseases, providing additional scale in treatments, but not providing a generalizable solution to the vast diversity of rare diseases^153^. Other more recent work has shown a novel prime-editing-installed suppressor tRNA approach for disease-agnostic gene editing that could theoretically be applied to conditions caused by nonsense mutations, which make up a significant fraction of genetic human diseases^154–156^. However, with the ability to generate active LSRs upon a specified attachment site used as prompt at inference time, we see that our aiPGI approach has the potential to scale even further in breadth and programmability of applications across cell and gene therapies.

For patients, this scalability is evidenced by the fact that EDEN generated active recombinases for all tested disease-associated loci and potential safe harbor sites, suggesting that thousands of currently intractable genomic targets are now within reach. Furthermore, this capability could significantly improve cell therapies by enabling the predictable insertion of large, multi-component genetic circuits into safe harbors, unlocking the sophisticated cellular logic required to tackle complex cancers and autoimmune diseases. This represents the beginning of a potentially transformative shift toward ‘mutation-agnostic’ medicines, where a single therapeutic product could treat heterogeneous genetic diseases, offering a potentially safer, one-time cure that overcomes the payload and genotoxicity limitations of current viral or nuclease-based editing.

On top of this, we show that the same model can design antimicrobial peptides (AMPs) with high prospective hit rates - 97% of tested AMPs showed activity, with top candidates achieving single-digit micromolar potency against critical-priority multidrug-resistant pathogens. To our knowledge, this is the first time a nucleotide-based foundation model has been used for the design of antibiotic peptides. Crucially, we demonstrate that this process is programmable: by prompting the model with pathogen-specific control tags, we steered generation toward specific targets, increasing the yield of high-confidence candidates against *E. coli* by approximately 16-fold compared to unconditional generation.

For patients facing the growing threat of antimicrobial resistance, this suggests a potential future capability where, in response to a resistant outbreak or a patient with a refractory infection like *Acinetobacter baumannii*, we can rapidly’dial in’ the specific target pathogen to generate bespoke, structurally novel antibiotic candidates on demand.

Finally, we extended the model’s capabilities beyond the organismal boundary to systems-level engineering. EDEN generated a gigabase-scale synthetic microbiome *in silico* that retained metabolic pathway completeness and complex host-phage relationships. This capability could open up the potential to engineer stable, multi-species consortia capable of correcting the dysbiosis underlying complex metabolic and immunological disorders.

Collectively, we believe that these results could represent an inflection point in generative biology. It is rare for a single foundation model to demonstrate, with robust wet-lab validation, the ability to design candidate therapeutic molecules across biochemically distinct regimes: By successfully spanning the small, amphipathic structures of antimicrobial peptides, the complex, multi-domain architecture of DNA-editing enzymes, and the gigabase-scale, metabolic logic of synthetic microbiomes, EDEN demonstrates that it has moved beyond narrow, task-specific optimization and is emerging as a versatile tool for generating effective therapeutic candidates across distinct modalities and disease areas.

We note that for therapeutic applications, the EDEN designed candidates will require continued validation in relevant human cells, and be profiled for site specificity with downstream optimisation through reinforcement learning or otherwise in order to increase activity and increase specificity. However, the transition from stochastic screening to predictable, prompt-based generation in response to therapeutically-relevant queries represents a meaningful shift in how we approach the engineering of biological systems.

### Toward a unified paradigm for predictable therapeutic design

When discussing the wider implications of biological foundation models targeted towards therapeutic design tasks, it is crucial to consider the potential of such models to address the most pressing issues in modern medicine. Modern day healthcare systems face a convergence of serious challenges: first and foremost the explosion of overall healthcare cost and the prohibitive expense of individual therapeutic discovery and development campaigns, occurring alongside a noticeable decline in pharmaceutical R&D efficiency known as Eroom’s law^157–160^. Simultaneously humanity faces significant biological threats, typified by the rise of antimicrobial resistance across a range of critical pathogens,^102^ and the systemic, growing burdens of cancer, genetic disorders, and autoimmune disease.

Given these harsh economic realities and biological threats, we argue that a scalable solution lies in the prospect of building unified AI systems that can design effective therapeutic candidates across modalities and molecule types, in a disease-agnostic, on-demand, and personalisable manner. To achieve this at scale, we return to our central hypothesis: that progress towards true programmable biology will require expanding the training datasets of generative models to include increasingly large quantities of diverse evolutionary data, far beyond the constraints of current publicly available resources.

Here, with our work on EDEN, we believe we have presented an early step that points towards that vision, with a single model demonstrating experimentally validated design capabilities across diverse modalities, molecule types, and disease areas, from small peptides to multiple complex gene insertion systems.

By demonstrating that state-of-the-art designs of therapeutic candidate molecules can be achieved using evolutionary priors alone, without human or clinical data in pre-training, these results support a shift in how biological foundation models can be constructed and applied. Crucially, this shift is enabled by overcoming a crucial data limitation that constrains models trained on public sequence data. While public databases face diminishing returns in high-quality, non-redundant diversity, BaseData’s purpose-built supply chain expands access to evolutionary & valuable cross-species sequence data at a scale supporting continuous model improvement and application range. In this respect, just as language models leveraged the vastness of the web to learn linguistic structure, biological models may increasingly leverage large-scale evolutionary data to learn and apply transferable biological principles towards increasingly complex and more therapeutically-aligned design tasks.

This suggests that a possible route to general-purpose biological intelligence lies not in generating massive amounts of clinical data, but in a hybrid approach: using large, scalable evolutionary datasets to learn universal design principles, which then act as a robust foundation for fine-tuning on smaller, high-value clinical datasets. Ultimately, we project that it will be this combination of billions of years of evolutionary data with specific therapeutic records that offers a potential scaling-driven path to making therapeutic design a predictable engineering discipline.

## Biosafety

The development and deployment of the EDEN generative foundation model family has been guided by rigorous biosafety and biosecurity considerations. All major applications of EDEN – from *de novo* recombinase engineering for gene insertion, to antimicrobial peptide generation and synthetic microbiome design – were accompanied by proactive risk considerations and oversight. Training data was stringently curated to exclude potentially hazardous sequences, notably filtering out all known eukaryotic viral genomes to prevent the inadvertent generation of pathogenic elements. Dual-use concerns have been carefully evaluated, particularly in relation to EDEN’s capacity to design genome-editing proteins and novel antimicrobials. For example, the World Health Organization’s *2022 Global Guidance Framework for the Responsible Use of Life Sciences* emphasizes comprehensive biorisk management spanning laboratory biosafety, biosecurity, and oversight of dual-use research and the U.S. National Science Advisory Board for Biosecurity (NSABB, 2023) has similarly called for integrated oversight of life science research with potential biosecurity risks^161,162^. EDEN’s development and evaluations adheres to all these principles. All laboratory work (such as validating EDEN-designed recombinases and peptides) was conducted under appropriate containment and institutional oversight, in accordance with NIH Guidelines for recombinant DNA research and institutional biosafety committee (IBC) review^162^.

### Stakeholder Best Practices for Equitable Data

All data used in EDEN pre-training has been collected with Prior Informed Consent (PIC), Material Transfer Agreements (MTA) and Mutually Agreed Terms (MAT) through Access and Benefit-Sharing (ABS) and knowledge-sharing partnerships that Basecamp Research has established across five continents. This means that each of the 9.7 trillion tokens used in EDEN pretraining can be traced back to stakeholder consent. Sample access is facilitated through 208 country-specific permits and 10 Access and Benefit-Sharing (ABS) collaboration agreements, which cover 28 countries in total, each explicitly defining the permission for the commercialisation of digital sequencing information (DSI) and describing the intended uses of the data - including downstream model training and other commercial applications - prior to collection. This approach ensures that consent is informed, freely given, and that the data is equitably sourced. Basecamp Research’s database provides full traceability of consent and permissions from the point of collection to downstream utilization, ensuring regulatory compliance and enabling the appropriate distribution of royalties back to Genetic Resource (GR) providers.

## Acknowledgements

We are grateful to the collaboration laboratories which have helped with the collection of samples for BaseData that was used for the training of this model, including: British Antarctic Survey (collaboration led by Professor Melody Clark), Heritage Malta, Malta (Professor Timmy Gambin), CENIBiot, a laboratory of CeNAT-CONARE, Costa Rica (Diego Hidalgo Badilla, Darling Mora, Professor Max Chavarría and the former director Dr. Randall Loaiza Montoya), Balaton Research Limnological Institute, Hungary (Dr. Kálmán Tapolczai), Wallowa Resources, USA (Nils Christoffersen), AJESH International, Cameroon (Harrison Ajebe Nnoko), Communities of Cameroon (Ongue Yassoukou, Nkom, Ngompem, and Bibema) and University of Buea (Dr. Eric Fokam). We further acknowledge our more recent collaborators who additionally continue to contribute to BaseData: the Scripps Oceanographic Institute, USA (Alexandra Hangsterfer), the Government of the Republic of Malawi, acting through Department of National Parks and Wildlife (DNPW), the Malawi University of Science and Technology (Dr Pedros Chigwechokha) and Codebreaker Bioscience, Chile (Alejandro Bisquertt).

Cesar de la Fuente-Nunez holds a Presidential Professorship at the University of Pennsylvania, and acknowledges funding from the Defense Threat Reduction Agency (DTRA; HDTRA1-21-1-0014, and HDTRA1-23-1-0001).

For supporting this work, we want to thank Saif Ur-Rehman, Richard de Napoli, James Ashton, Phoebe Oldach, Ineke Knot, Marlon Clark, Silvano Ulivi, Nadine Greenhalgh, Bupe Mwambingu, Leif Christofferson, Luis Enrique Garcia Rivera, Jonathan Curry, Konstantinos Andrikopoulos, Nick Coleman, D’Arcy Doran, Angus MacPherson, Victor Zhukov, Rene Schönfelder, and Ilya Burkov.

For insightful discussions, helpful feedback, and input, we want to thank in particular David R. Liu, Rohit Sharma, Omar Abudayyhe, as well as the de la Fuente Lab.

## Conflicts of Interest

Some of the authors are current or former employees of Basecamp Research Ltd. and its wholly-owned subsidiary Basecamp Research US Inc. and may hold shares in either. BaseData™, BaseFold™, PGI™, aiPGI™, EDEN-GLM™, EDEN-LSR™ are brand names and technologies of Basecamp Research Ltd. Cesar de la Fuente-Nunez is a co-founder of, and scientific advisor, to Peptaris, Inc., provides consulting services to Invaio Sciences, and is a member of the Scientific Advisory Boards of Nowture S.L., Peptidus, European Biotech Venture Builder, the Peptide Drug Hunting Consortium (PDHC), ePhective Therapeutics, Inc., and Phare Bio.

## IP Statement

Some of the authors that are employees of Basecamp Research Ltd and its wholly-owned subsidiary Basecamp Research US Inc. are inventors or co-inventors on pending patent applications that encompass innovations related to disclosed subject matter in this manuscript.

## Methods

### BaseData and OG2 curation

We applied stringent filtering criteria to retain only high-quality contigs. We first required contigs to be longer than 2 kb and to exhibit a predicted gene density greater than 20%. In addition, contigs were required to have a minimum mean sequencing depth of at least 4X. Low-complexity contigs shorter than 10 kb and containing more than 50% low-complexity sequence, as quantified with DUSTmasker (v2.15.0), were removed^163^. Finally, contigs with hits to known eukaryotic viruses, as identified by GeNomad (v1.7.6) annotation, were excluded from the final dataset^144^. For the 100M, 1B, and 7B parameter models trained on BaseData, we fixed the random seed and used an identical data split across all training runs, training each model on up to 350B tokens, whereas the 28B parameter model was trained on the entirety of BaseData. We used the metagenomic portion of the OG2 dataset for the training of the 100M, 1B, and 7B models and followed the same training procedure as we did for BaseData by fixing a random seed and using an identical data split across training runs.

### EDEN architecture

The EDEN model family adopts a decoder-only Transformer architecture. The largest configuration uses up to 48 layers with a hidden dimension of 6,144 and 48 attention heads. Across all model scales (100M, 1B, 7B, and 28B parameters), we use 8 key-value heads, SwiGLU activation, a vocabulary size of 512, and set the RoPE base frequency hyperparameter to 500,000. We use the Llama 3.1 implementation in BioNemo (using NeMo 2.0 and Megatron-LM^124^). (Table 2).

**Table 2:** Overview of the key hyperparameters of EDEN. We display settings for 100M, 1B and 7B and 28B models.

| Hyperparameter | 100M | 1B | 7B | 28B |
| --- | --- | --- | --- | --- |
| Layers | 12 | 16 | 32 | 48 |
| Model Dimension | 512 | 2048 | 4096 | 6144 |
| FFN Dimension | 2048 | 8192 | 14336 | 26368 |
| Attention Heads | 8 | 16 | 32 | 48 |

### Pre-training

EDEN-28B was pre-trained using the AdamW optimizer (β₁ = 0.9, β₂ = 0.95, ε = 1 × 10⁻⁸, weight decay = 0.01) with global norm gradient clipping set to 1.0. The learning rate schedule followed cosine decay with an initial linear warmup for the first 2,500 steps, peaking at 3 × 10⁻⁵ and annealing to a minimum of 6 × 10⁻⁷ over a total of 640,920 optimizer steps. Training was performed using bfloat16 mixed precision, with FP8 hybrid mode enabled. A global batch size of 2,016 sequences was used (micro-batch size of 1, gradient accumulation 4, data parallelism 504), each with a sequence length of 8,192, resulting in approximately 16.52 million tokens processed per update. The parallel training setup consisted of tensor parallelism = 2, with sequence parallelism enabled, utilizing 1,008 GPUs deployed across 126 nodes (data parallel = 504).

In accordance with the stabilization method proposed by Takase and Kiyono (2025), the embedding layer was initialized from a normal distribution, Ɲ(0,1.0)^164^, while the remaining model parameters used standard Megatron initialization. For efficiency and robustness, the training process included overlapped gradient reduction, asynchronous checkpointing, and preemption support; fp32 residual connections were disabled. Training remained stable throughout and, although occasional loss spikes were detected, no intervention was required to correct for model training divergence.

### Dataloading

EDEN tokenizes DNA sequences at single-nucleotide resolution, using a byte-level tokenizer with an effective vocabulary of four tokens, one per base, from a total vocabulary of 512 characters. We employed a sliding window dataloader to process genomic sequences of arbitrary length. Each sequence was partitioned into overlapping windows of 8,192 tokens with a stride of 7,992 tokens, resulting in 200 bp overlap between consecutive windows. This overlap preserves local context at window boundaries. Each window is formatted as: BOS token, followed by optional control tags, a SEP token, the sequence, and an EOS token.

### Compute

EDEN-28B was trained on 1,008 H200 GPUs, each running at 700W TDP with 141GB HBM3e memory. Each server is equipped with eight GPUs, within a server the GPUs are connected with NVLink, servers are interconnected to each other with NVIDIA Quantum-2 InfiniBand at 400Gb/s. Training jobs are scheduled with the Kubeflow Pytorch Operator on Kubernetes. For storage we used the Nebius Shared Filesystem storage fabric, backed by all-flash NVMe drives and served over NFS. For Networking EDEN used a NVIDIA Quantum-2 InfiniBand fabric at 400 Gb/s between nodes

### Inference

EDEN inference was performed using two inference backends: the inference pipeline from the NVIDIA BioNeMo framework (v2.6.3)^165^ and the vLLM OpenAI-compatible server (v0.11.0)^142^. For vLLM, optimized implementations for Llama 3.1 were used, with FlashInfer (v0.4.1)^166^ enabled to support efficient attention computation. vLLM was utilized as an alternative inference engine due to its optimisations for language-model inference and ease of deployment. Evo2 inference was performed using the Evo2 NVIDIA NIM (NVIDIA Inference Microservices) (v2.1.0)^167^ following the deployment guidelines from NVIDIA. EDEN and Evo2 inference was performed on NVIDIA H200 GPUs.

### Scaling laws

To characterise scaling behavior across model capacities, we trained the EDEN models at three parameter scales (100M, 1B and 7B). For comparability, training used randomly sampled subsets matched for total nucleotide budget (350 billion nucleotides per dataset). Cumulative training compute was estimated from empirical throughput statistics. Raw floating-point operations (FLOPs) were derived as the product of total training steps, measured GPU throughput, and step duration. Given the variable sequence lengths between datasets, we normalized raw compute to effective FLOPs to account for padding overhead in the absence of sequence packing. We empirically determined the fraction of loss-contributing tokens by randomly sampling five global batches from each dataset (approximately 84 million tokens). This correction ensures that scaling metrics reflect compute allocated strictly to valid biological signals.

Model performance was assessed via test perplexity (on a held-out test set from both datasets) at checkpoints matched for total nucleotide exposure. To quantify data efficiency, we modeled the relationship between perplexity and effective compute as a power law. The scaling parameters exponent was estimated via linear regression in log-log space. We observed fitted scaling exponents of 𝑏≈0.058 for BaseData and 𝑏≈0.047 for OG2. Within the range of model sizes we tested, the BaseData models exhibit a steeper scaling exponent and lower perplexity at high FLOPs/model size than the OG2 models. This provides evidence that BaseData is effectively “higher quality” in the scaling-law sense: as compute and capacity increase, perplexity decreases faster for BaseData than for OG2.

### BaseData genomic uniform manifold approximation and projection (UMAP)

K-mer frequencies (k=4-6) were calculated for all BaseData contigs using Jellyfish v2.3.1^168^. A maximum of 10,000 contigs were selected from each sample and a metadata-based scaling factor was then applied using the Lineage 3 MGnify biome for the sample to visually separate contigs with different biome metadata^82^. A UMAP of Euclidean distances for the scaled k-mer frequencies was calculated using the UMAP function of umap-learn v0.5.9^169^ with parameters n_neighbors=20, min_dist=0.1 using a random subset of 10,000,000 contigs. The UMAP embeddings were plotted using Datashader v0.18.2^170^.

### Gene autocompletion

Prompt sets were constructed from conserved genes *recA* from *Bacillus subtilis*, *secY* from *Streptomyces coelicolor*, and *ftsZ* from *Escherichia coli* using 20% of each native DNA sequence as the prompt. Sequence generation produced 1500 bp outputs with top k 4 and temperature 0.9. The generated DNA sequences were translated into protein sequences and percentage identity to the corresponding natural proteins was calculated using global alignment implemented in Biopython. Translated proteins were folded using ESMFold and TM-score against the native protein was calculated using USalign, normalising to the length of the reference structure^66,171^.

Additionally, prompt sets comprising 30% of large serine recombinase sequences from BaseData were generated and used to produce 2500 bp sequences. The prompts were prepended to the generated DNA sequences, which were then translated into protein sequences, and ESM embeddings were computed and projected using UMAP.

### Operon autocompletion

Prompt sets for trp operon and modABC operon from E. coli K-12 (NC_000913.3) were generated using the full coding sequences of each gene in the operon, including reverse complement sequences for bi-directional generation. For each prompt, sequences of length 2500 nt were generated using generation parameters top k 4 and temperature 0.9. After gene prediction using pyrodigal (version 3.6.0), predicted proteins were folded using ESMFold and TM-score against the native protein was calculated using USalign, normalising to the length of the reference structure^135,171,172^. Native and generated protein structures were subsequently visualized using PyMOL.

### Evaluation of long context generations

A prompt set targeting the bacterial S10 operon was curated using ten publicly available NCBI RefSeq assemblies (Table 3). The start coordinates of the operon in each reference was determined by annotating the assemblies with Bakta v1.8.1^173^ and light database v2025-02-24. Samtools v1.22.1^174^ was used to extract the first 1000bp of the operon in each reference as a prompt set, and 14,000bp downstream of the start of each was extracted as a point of reference. EDEN-28b and evo2-40b-8k were prompted with each of the ten sequences, specifying 100 generations per prompt at 13,000 tokens with top-k=4, top-p=1 and temperature=0.9. Coding densities were calculated by predicting ORFs in each generated sequence with Pyrodigal v3.6.3 with option-p meta^172^. The proportion of nucleotides occurring within ORFs was calculated across a sliding window of 100bp across each generated sequence. The percentages of unmasked bases across each 100bp window was calculated as 100 minus the percentage of bases masked using pydustmasker v1.0.3 with default light database v2025-02-24 and arrow plots of the operons created using lovis4u v0.1.6^173,175^. Predicted proteins were folded using ESMFold and TM-score against the native protein was calculated using USalign, normalising to the length of the reference structure^171^ and visualized using PyMOL.

**Table 3:**
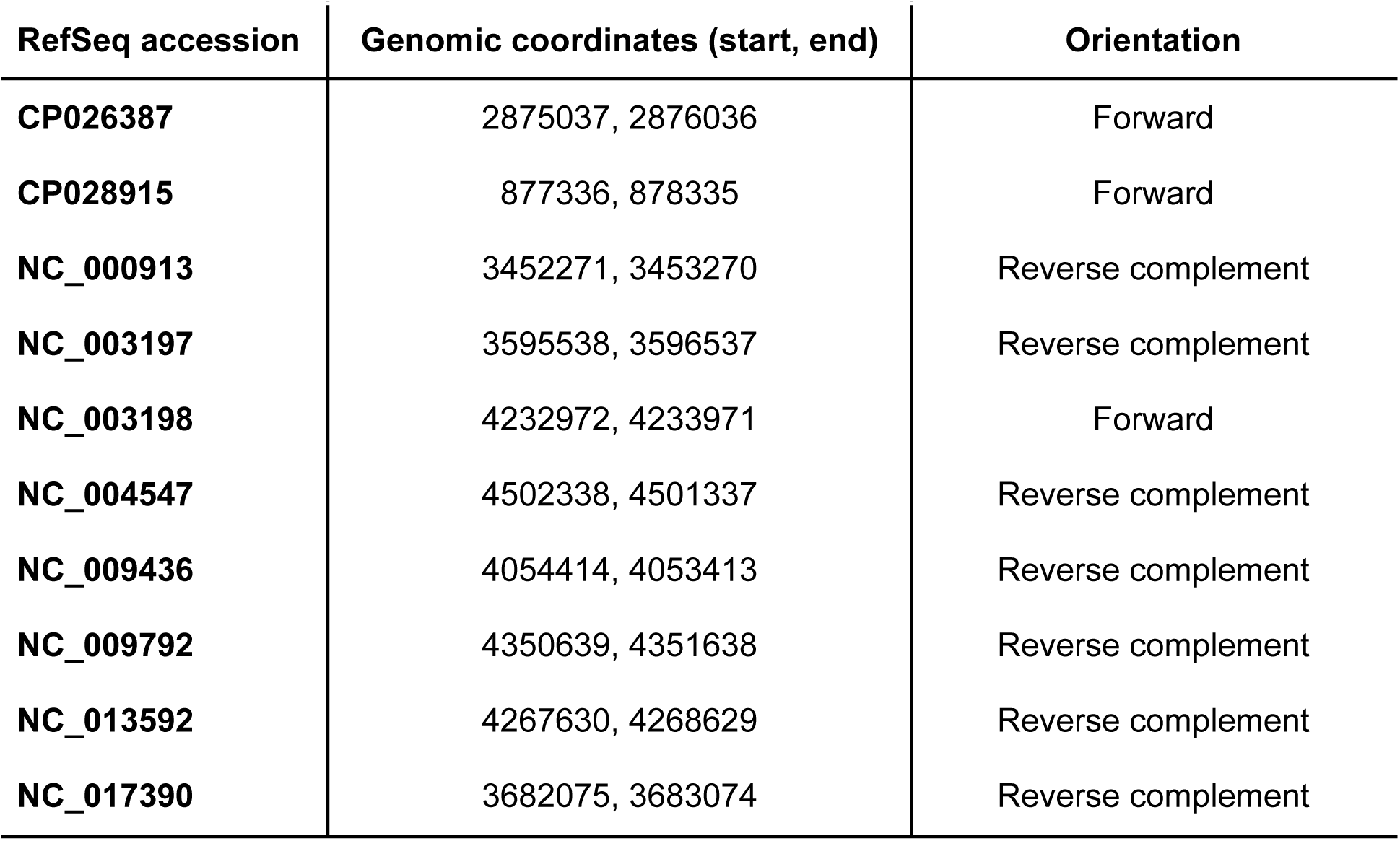
Public prompts used in the long-context generations.

### Protein mutational effect prediction

We used published deep mutational scanning (DMS) datasets to benchmark EDEN against RNA and DNA models in their ability to predict the functional consequences of mutations. Specifically, we used the prokaryotic datasets curated in RNAGym^176^ to evaluate mutational effect prediction performance for coding genes. When scoring sequences with EDEN-28B, a BOS token was prepended to the start of each sequence. In addition to the results reported in RNAGym, we evaluated Evo2-40B on all datasets, with an EOS token prepended to each sequence.

### EDEN fine-tuning with LSRs

Prior to training, paired LSR-attachment site sequences were clustered at a 70% sequence identity threshold using MMseqs2, with cluster singletons held out as an independent validation set. EDEN was fine-tuned using a batch size of 32 and the model checkpoint with the lowest cross entropy loss on the validation set was retained for evaluation. For fine-tuning, clusters were sampled uniformly, irrespective of their size, and individual LSR-attachment site pairs were selected from each cluster. Training employed the Adam optimizer. The learning rate was increased from zero to a maximum of 9e-6 and decayed according to a cosine annealing schedule. The model was trained to optimise the cross entropy loss over the next token distribution.

### EDEN synthetic LSR generation

LSR generation was prompted with either full attBoP′ (60 nucleotides) or attB-half (30 nt) context from active LSR-attachment site pairs. Generated DNA sequences were first translated into amino acid sequences with Biopython, and InterProScan (v5.75-106.0) was run to assign putative LSRs by the sequential presence of PF00239 (*Resolvase, N terminal domain*), PF07508 (*Recombinase*), and PF13408 (*Recombinase zinc beta ribbon domain*). Synthetic LSRs with fewer than 700aa, high sequence complexity (computed by tantan and sequence entropy), and at least 50% Levenshtein sequence identity to the wildtype LSR for a given attachment site were selected as candidates for wet lab experimentation^177^. For determination of genomic sites in putative safe harbors, we used previous literature information^178^

### LSR production and quantitative PCR recombination assay

Double-stranded DNA (dsDNA) substrates were synthesized in arrayed format (Twist Bioscience). Each substrate contained, in linear order, an attB site, a T7 promoter, the recombinase ORF, and an attP site. Large serine recombinase (LSR) protein was expressed from the dsDNA template using PURExpress® In Vitro Protein Synthesis Kit (NEB catalog #E6800). Unless otherwise indicated, the final concentration of the LSR template in IVTT reactions was 10 nM.

Recombination was assayed by self-circularization of the dsDNA template mediated by intramolecular attB–attP recombination. Circularization juxtaposes primer binding sites that are oriented away from one another on the linear template but face each other following recombination. Reaction products were quantified by quantitative PCR (qPCR) (NEB catalog #M3003) using primers (IDT or Azenta) specific to the recombined junction.

All qPCR assays were performed in technical duplicate, alongside a single no-IVTT dsDNA negative control lacking transcription–translation reagents. Quantification was based on relative cycle threshold (Cq) values using a fixed reference point applied uniformly across all reactions, such that larger ΔCq values reflected greater accumulation of recombined product. The threshold was selected such that all positive samples had a one-sided tail probability of approximately p=0.01 under a Gaussian assumption, with ΔCq values from negative-control measurements used to define the null distribution.

### Multiplexed quantitative PCR recombination assay

Nine pseudosite substrates were designed, each consisting of an attH sequence, a spacer region, and an attP sequence (Twist Bioscience). One of three unique primer pair binding site and TaqMan probe binding site combinations was embedded within the spacer region of each substrate to enable multiplexed detection of circularized recombination products. Each combination/assay was assigned a unique fluorophore in combination with a ZEN™/TAO™ internal quencher and/or 3’ Iowa Black™ FQ/RQ quencher (IDT). A wild-type attB–attP substrate (under a unique primer/probe set and fluorescent channel) was included as an internal reference in all reactions.

Substrates were pooled into three multiplex reactions, each containing three pseudosite substrates and the wild-type substrate for a total of four substrates per pool. LSR-coding gene fragments (Twist Bioscience) were incubated with each substrate pool (3) in separate reactions for 15 minutes at 37°C using PURExpress® In Vitro Protein Synthesis Kit (NEB catalog #E6800) followed by 1:20 dilution in ultra-pure water before multiplexed quantitative PCR alongside no-IVTT pooled-substrate dsDNA negative controls (3; one per pool) using PrimeTime™ Gene Expression Master Mix (IDT catalog #1055772) on a QuantStudio™ 6 Pro Real-Time PCR System (ThermoFisher). Primer concentrations were empirically adjusted to balance amplification maxima between wild-type and pseudosite substrates.

Recombination efficiency for each substrate was quantified based on Cq values obtained from the corresponding fluorescence channel. Quantification was based on relative cycle threshold (Cq) values using a fixed reference point applied uniformly across all reactions. Quantification was based on relative cycle threshold (Cq) values using a fixed reference point applied uniformly across all reactions, such that larger ΔCq values reflected greater accumulation of recombined product. The threshold was selected such that all positive samples had a one-sided tail probability of approximately p=0.01 under a Gaussian assumption, with ΔCq values from LSR reactions with low/no detectable amplification (presumed negative) used to define the null distribution.

### IVTT-based cryptic sequence recombination assay

The pseudosite recombination discovery assay was adapted from the previously described Cryptic-Seq protocol ^95^. Recombinase protein was produced by IVTT from linear dsDNA templates encoding the corresponding ORFs using PURExpress® In Vitro Protein Synthesis Kit (NEB catalog #E6800). Human male (XY) genomic DNA (Millipore Sigma catalog #70572) was prepared as an integration target background via enzymatic fragmentation and next-generation sequencing adapter ligation (custom; P7-only 5’ overhang with UMI) using NEBNext® Ultra™ II FS DNA Library Prep Kit for Illumina (NEB catalog #E7805).

Linear dsDNA donor fragments containing candidate cryptic recombination sites were incubated with NGS-adaptor-ligated gDNA libraries and recombinase protein IVTT product at 37°C overnight. Libraries of integrated donor DNA junctions were amplified via nested target-enrichment PCR and sequenced on an Illumina MiSeq i100+ platform using paired-end sequencing. Sequencing data were processed using a custom bioinformatics pipeline designed to detect split reads spanning junctions between donor fragments and human genomic DNA; reads containing one segment mapping to the donor fragment and a second segment mapping to the human genome were classified as recombination events and used to identify cryptic recombination sites.

### Human cell activity assay

K562 cells were maintained in RPMI 1640 Medium (Gibco 11875093) with 10% FBS (Gibco A3840201) at 37C 5% CO2. 100,000 cells per condition were electroplated using the Neon™ NxT Electroporation System in a 10uL total volume to introduce 2ug mRNA encoding the LSR (PRT075) and 2ug cargo plasmid containing the corresponding attP sequence. After 5 days in culture, genomic DNA was extracted using QuickExtract (QE09050) according to the manufacturer protocol. dPCR was run on 10ng of DNA for each condition (QuantStudio Absolute Q) to quantify the integration at a pseudosite on chromosome 7. RNaseP was used as a housekeeping gene for normalization. T cells were isolated from a Leukopak using negative selection using EasySep™ Human T Cell Isolation Kit (STEMCELL 17951), banked and stored in LN2. At day-3, cells were thawed into ImmunoCult™-XF T Cell Expansion Medium (STEMCELL 10981) supplemented with 10ng/mL Human Recombinant IL-2 (STEMCELL 78145) and activated with Dynabeads™ Human T-Activator CD3/CD28 for T Cell Expansion and Activation (11161D) for 3 days. On D0, the Dynabeads were removed and 100,000 cells were electroplated using the Neon™ NxT Electroporation System in a 10uL total volume to introduce 1.5ug mRNA encoding the LSRs (WT LSR and 20 EDEN generated LSR’s) and 1ug cargo plasmid with EF1a driving the expression of a CD19 CAR and containing the corresponding attP sequence. 3 days after transfection, cells were stained with LIVE/DEAD™ Fixable Violet Dead Cell Stain (Thermo L34964) and PE-Labeled Human CD19 and analyzed on the Attune NxT Flow Cytometer. Percent integration in live cells was quantified using a cargo only control as the negative gate.

### EDEN fine-tuning with bridge recombinase (BR) systems

EDEN-28B was fine-tuned using a dataset of putative Bridge Recombinase (BR) systems, which included both the bDNA and the bridge recombinase protein sequences. This dataset, consisting of 6,183,225 sequences, specifically focused on ORFs containing the PF01548 (*DEDD_Tnp_IS110*) and PF02371 (*Transposase_20*) protein domains, along with 1 kilobase of sequence both upstream and downstream (Fig. 4B).

To prevent data leakage, amino acid sequences were clustered at 80% identity using mmseqs2 (v15.6f452)^179^. The resulting clusters were then split into training, validation, and test sets at a ratio of 80%, 10%, and 10%, respectively. Fine-tuning was performed using the bionemo codebase over 21500 steps. Model evaluation utilized a set of 239 BR sequences, comprising a mix of 72 publicly available^129^ and 167 proprietary sequences. The sequences comprising the benchmark were not utilized in the fine-tuning process. For each sequence in the benchmark, four prompts were generated, corresponding to 25%, 50%, 75%, and 100% of the bDNA sequence. One hundred sequences were generated for each prompt condition. Valid generations were selected using the same parameters defined in the base model section.

### Computational evaluation of generated BRs

Generation (for both zero-shot and fine-tuned models) was prompted using either 25%, 50%, 75% or 100% of the bridge guide upstream of the recombinase coding sequence. Generated DNA sequences were first translated into ORFs with prodigal (metagenomics mode), proteins between 300-500aa and starting with a methionine were retained for downstream analysis. InterProScan was run to assign ORFs as putative BRs by presence of PF01548 (*DEDD_Tnp_IS110*) and PF02371 (*Transposase_20*) domains. In addition, those with DEDD residues and the conserved serine residue corresponding to position 241 in IS621 underwent structure prediction using BaseFold^180^. Predicted structures bearing a pLDDT score between 70-90, and those sharing structural similarity (i.e. RMSD ≤ 3 over 300aa) to IS621 or IS622 are nominated for *in vitro* validation. With the exception of the 100% guide prompt, the generation of guide fragments necessitates evaluation of the completion of the guide via CMsearch based on a covariate model containing guide sequences of IS621 and related public and BaseData sequences, and matching systems with an e-value of < 0.01 were considered for manual detection of putative target and donor-binding and handshake residues^129^.

### Laboratory validation and evaluation of activity in generated BR systems

Double-stranded DNA (dsDNA) fragments of bridge recombinases, bridge RNA and cargo were synthesized in arrayed format from IDT or Twist Biosciences. The bridge recombinase dsDNA fragment contained in linear order a T7 promoter, 5’ UTR, recombinase ORF and 3’ UTR. The bridge RNA dsDNA contained a T7 promoter, the bridge RNA sequence containing target and donor loops. The cargo fragment contained target and donor sequences and outward facing. Primers were synthesized by IDT to amplify each dsDNA fragment. Amplifications were carried out using Platinum™ SuperFi™ PCR Master Mix (Invitrogen, Catalog# 12358050) and PCR product was purified using AMPure XP Beads (Beckman Coulter, Catalog# A63881). Concentrations were measured using Qubit DNA BR kit (Thermo Fisher Scientific, Catalog# Q33266). IVTT reaction was carried out using PURExpress® In Vitro Protein Synthesis Kit (New England Biolabs, Catalog# E6800) with dsDNA fragment of bridge recombinase and bridge RNA. dsDNA cargo fragment was incubated with the IVTT product, followed by heat inactivation. For quantification of circularisation, outward facing primers on the cargo dsDNA fragment that detect recombined circular products (similar to the LSR quantitative PCR recombination assay) were used. qPCR for samples were set up using Luna qPCR Master Mix (New England Biolabs, Catalog# M3003) in technical replicates alongside a single no-IVTT dsDNA negative control lacking transcription translation reagents. Cq values were assessed to determine abundance of circular products. ΔCq values were calculated based on their respective controls and used to determine activity of the recombinase system, where higher ΔCq value represents higher accumulation of recombination products.

### Antimicrobial peptide discovery

ORFs/smORFs were mined from assemblies in the Basecamp Research BaseData^1^ database. After filtering for candidates <=50aa, APEX-pathogen^133^ (version taken Apr 22, 2025) was run on candidate ORF/peptide sequences. For each peptide, we calculate the median predicted MIC (µM) across strains as a holistic metric for classifying antibiotic activity. A threshold of 64 (µM) was used to filter candidates. This subset of candidates were compared against the DRAMP database (v4.0)^134^ using a StripedSmithWaterman local alignment and BLOSUM50 scoring matrix and a 0.7 similarity cutoff for novel classification. Additional annotation for taxonomy using the underlying contig the candidate ORF is on was done using kraken2(v2.1.3)^181^ taxonomic classifier.

From the filtered AMP discovery candidates, mmseqs2 (v15.6f452)^179^ cluster was run with sequence ID and coverage cutoffs of 0.9. A custom script was written to return the best representative for each cluster according to the median predicted MIC value. Representatives were broken down to three tiers based on predicted median MIC cutoffs. Candidates were chosen from those with a median MIC cutoff of 32 (µM). These candidates were then filtered and selected based on sequence composition and likelihood of synthesis success.

### EDEN synthetic antimicrobial peptides generation

Two fine-tuning datasets were curated from BaseData based on predicted MIC values using apex-pathogen (version taken Apr 22, 2025)^133^. The first dataset comprised ORFs with a predicted median MIC below 64 µM and included 8 kb of surrounding genomic context. The second dataset comprised ORFs that have a predicted MIC of less than 64 µM for any pathogen, and AMP-specific start and end tags and tags pertaining to the predicted pathogens it targets were added to the sequences. Each dataset was split into training, validation, and test sets using an 80%, 10%, and 10% partition, respectively. Using these splits, two models were derived from the EDEN-28B base model via full-parameter fine-tuning (Figure 5A): one trained on the 8 kb context window dataset and the other trained on the pathogen-tagged ORF dataset.

Generation of antimicrobial peptides was conducted using the fine-tuned 8Kb context model and the fine-tuned pathogen tag model. The prompt strategy involving genomic context sequences were constructed using either 300 bp or 1 kb of upstream sequence, with or without including a start codon. Source sequences were drawn from three sets: AMP-containing contigs chosen at random curated from BaseData, contigs containing AMPs with a predicted MIC below 32 µM, and representatives from the top 20 sequence clusters. Sequence generation was performed via a vLLM code inference endpoint, producing 100 sequences per prompt. Generation using the fine-tuned tag model was conducted by prompting on the specific pathogen or start tag using a custom script leveraging the bionemo framework.

Generated sequences were translated using biopython, filtered to only include sequences under 50 amino acids, and apex-pathogen was used to predict the per-strain MIC. Peptides with a predicted MIC of ≤ 32 µM for at least one strain were retained for downstream analyses. Physicochemical properties (charge, isoelectric point, hydrophobicity, and hydrophobic moment) of the generated AMPs were calculated using modlAMP (v 4.3.2)^182^. To compare EDEN-generated AMPs with reference AMP collections, sequence similarity scores were computed against entries in the DRAMP (v4.0) database^134^ and BaseData AMPs used to fine-tune the models. Sequences were aligned against the target database using the Smith-Waterman local alignment algorithm with the BLOSUM50 substitution matrix. The sequence similarity score between these two peptides was defined as the normalized alignment score as defined previously^110,112^. Additionally, peptides from DRAMP, BaseData, and EDEN were embedded with the ESM-2 650M model and projected into two dimensions using UMAP (cosine distance; n_neighbors = 15; min_dist = 0.1) for visualisation. Lab-tested peptides were matched to their embeddings and overlaid on the same projection.

Additional annotations for filtering of candidates for experimental validation included, charge (+2 to +8 net for 25 to 40-mers) and hydrophobicity of residue counts (35–60% hydrophobic/aliphatic residues), presence of aggregation patterns (>2 aromatics in a row, >4 hydrophobics in a 6-residue window, or highly repetitive triads) and similarity to training data. This set was then filtered with a length cutoff of <40aa. Candidates were additionally clustered at 80% using mmseqs2 (v15.6f452)^179^ Candidates that passed this initial filtering were then analysed for simple repeats and low complexity regions using tantan default parameters (version 51)^177^. Percent of masked regions were then calculated for each candidate sequence. Sequences over 20% repeat across the length were excluded. Focus on the predicted MIC scores for pathogens *A. baumannii* ATCC 19606*, E. coli* AIC221*, E. coli* AIC222 activity was prioritized as the predicted results for these pathogens show stronger predictive values for activity. A Smith-Waterman similarity score threshold of 0.7 was used, with only sequences scoring below this threshold advanced to wet-lab validation.

### Peptide synthesis and characterization

Peptides were synthesized on an automated peptide synthesizer (Symphony X, Gyros Protein Technologies) by standard 9-fluorenylmethyloxycarbonyl (Fmoc)-based solid-phase peptide synthesis (SPPS) on Fmoc-protected amino acid-Wang resins (100–200 mesh). In addition to preloaded resins, standard Fmoc-protected amino acids were employed for chain elongation. N,N-Dimethylformamide (DMF) was used as the primary solvent throughout synthesis. Stock solutions included: 500 mmol L^-1^ Fmoc-protected amino acids in DMF, a coupling mixture of HBTU (450 mmol L^-1^) and N-methylmorpholine (NMM, 900 mmol L^-1^) in DMF, and 20% (v/v) piperidine in DMF for Fmoc deprotection. After synthesis, peptides were deprotected and cleaved from the resin using a cleavage cocktail of trifluoroacetic (TFA)/triisopropylsilane (TIS)/dithiothreitol (DTT)/water (92.8% v/v, 1.1% v/v, 0.9% w/v, 4.8%, w/w) for 2.5 hours with stirring at room temperature. The resin was removed by vacuum filtration, and the peptide-containing solution was collected. Crude peptides were precipitated with cold diethyl ether and incubated for 20 min at-20 °C, pelleted by centrifugation, and washed once more with cold diethyl ether. The resulting pellets were dissolved in 0.1% (v/v) aqueous formic acid and incubated overnight at-20 °C, followed by lyophilization to obtain dried peptides.

For characterization, peptides were dried, reconstituted in 0.1% formic acid, and quantified spectrophotometrically. Peptide separations were performed on a Waters XBridge C_18_ column (4.6 × 50 mm, 3.5 µm, 120 Å) at room temperature using a conventional high-performance liquid chromatography (HPLC) system. Mobile phases were water with 0.1% formic acid (solvent A) and acetonitrile with 0.1% formic acid (solvent B). A linear gradient of 1–95% B over 7 min was applied at 1.5 mL min^-1^. UV detection was monitored at 220 nm. Eluates were analyzed on Waters SQ Detector 2 with electrospray ionization in positive mode. Full scan spectra were collected over m/z 100–2,000. Selected Ion Recording (SIR) was used for targeted peptides. Source conditions were capillary voltage 3.0 kV, cone voltage 25-40 V, source temperature 120 °C, and desolvation temperature 350 °C. Mass spectra were processed with MassLynx software. Observed peptide masses were compared with theoretical values, and quantitative analysis was based on integrated SIR peak areas.

### Bacterial strains and growth conditions

The bacterial panel utilized in this study consisted of the following pathogenic strains: *Acinetobacter baumannii* ATCC 19606; *A. baumannii* ATCC BAA-1605 (resistant to ceftazidime, gentamicin, ticarcillin, piperacillin, aztreonam, cefepime, ciprofloxacin, imipenem, and meropenem); *Escherichia coli* ATCC 11775; *E. coli* AIC221 [MG1655 phnE_2::FRT, polymyxin-sensitive control]; E. coli AIC222 [MG1655 pmrA53 phnE_2::FRT, polymyxin-resistant]; E. coli ATCC BAA-3170 (resistant to colistin and polymyxin B); *E. coli* K-12 BW25113; *Enterobacter cloacae* ATCC 13047; *Klebsiella pneumoniae* ATCC 13883; *K. pneumoniae* ATCC BAA-2342 (resistant to ertapenem and imipenem); *Pseudomonas aeruginosa* PAO1; *P. aeruginosa* PA14; *P. aeruginosa* ATCC BAA-3197 (resistant to fluoroquinolones, β-lactams, and carbapenems); *Salmonella enterica* ATCC 9150; *S. enterica* subsp. enterica Typhimurium ATCC 700720; *Bacillus subtilis* ATCC 23857; *Staphylococcus aureus* ATCC 12600; *S. aureus* ATCC BAA-1556 (methicillin-resistant); *Enterococcus faecalis* ATCC 700802 (vancomycin-resistant); and *Enterococcus faecium* ATCC 700221 (vancomycin-resistant). *P. aeruginosa* strains were propagated on *Pseudomonas* Isolation Agar, whereas all other species were maintained on Luria-Bertani (LB) agar and broth. For each assay, cultures were initiated from single colonies, incubated overnight at 37 °C, and subsequently diluted 1:100 into fresh medium to obtain cells in mid-logarithmic phase.

### AMP Minimal Inhibitory Concentration (MIC) determination

MIC values were established using the standard broth microdilution method in untreated 96-well plates. Test peptides were dissolved in sterile water and prepared as twofold serial dilutions ranging from 1 to 64 μmol L^-1^. Each dilution was combined at a 1:1 ratio with LB broth containing 4 × 10^6^ CFU mL^-1^ of the target bacterial strain. Plates were incubated at 37 °C for 24 h, and the MIC was defined as the lowest peptide concentration that completely inhibited visible bacterial growth. All experiments were conducted independently in triplicate.

### Synthetic metagenomes: fine-tuning, generation, and characterization

Characterization of biomes based on annotations was conducted on assembled contigs contained in BaseData. Digestive system specific annotations were identified and used as markers for sequence collection. GC and coding density were also used to characterize biomes. Biome labels are based on MGnify names and lineages^82,183^.

Before large scale sequence generation, EDEN was fine-tuned for 1.5 epochs on 160,885 assembled contigs from a single digestive-system microbiome sample in BaseData.For synthetic microbiome generation, we identified prompts from curated digestive system sequences which fell into three categories: sequences from the beginning of BaseData digestive system sample contigs, genes enriched in BaseData digestive system samples, and genes within 5kb neighborhoods known to co-occur with other conserved genes in BaseData digestive system samples. Source of sequence prompts were from BaseData. Each category of prompt contained between 150-200 carefully curated sequences between 1000 bp to the length of an entire gene to generate 10kb sequences. Generation was facilitated using a custom script leveraging the specific vLLM code inference endpoint.

Generated sequences of 10kb were initially filtered using length and low complexity criteria. This subset was then run through an internally developed custom annotation pipeline. This pipeline characterizes genes, proteins and pathways. The generated sequences were run through pyrodigal (3.6.0) to call ORFs^172^. Diamond (v2.1.14) was used to align ORFs against the kegg database (2011)^139^. InterProScan (v5.76-107.0local) was used for functional annotation of domains using PFAMs^184^. Skani (v0.3.0) was used to compare generated sequences with BaseData^185^. We aligned the source short reads from the sample used in fine tuning against all generated sequences using strobealign (v0.16.1)^186^. Viral and phage annotation was conducted using geNomad (v1.11.1)^144^.

A list of ORFs with KEGG annotations across all contigs were run through GSEApy (v1.1.10) package’s enrichr function using a custom gmt pathway database curated from BioCyc and annotated with KEGG ids^139,187,188^. The list of resulting enriched pathways were filtered to keep pathways with adjusted p-value <= 0.05.

For prophage generation, we leveraged our fine tuned model to run a second round of generations prompting it on the end of our generations from the first set that were categorized as a viral contig or as containing a pro-phage by genomad (v1.9.4, database v1.9)^144^. We then concatenated the original sequence and new sequence to obtain a sequence 21kb in length and ran this through genomad (v1.9.4, database v1.9) to obtain a 7kb pro-phage like sequence with viral score > 0.9 and taxonomy assigned to *Caudoviricetes* flanked by sequences from the host genome (*Streptomyces* on both ends) Sequences were then annotated with pharokka v1.8.2 and phold v1.1.0 to create consensus annotations, and visualisations of prophage candidates created using phold plot and lovis4u v0.1.6^146,147,175^.

### Taxonomic UMAP

BaseData samples were sub-sampled to contigs between 7-10kbp in length to match the length distribution of contigs in the synthetic metagenome. Taxonomic profiling of the synthetic metagenome and all filtered BaseData samples was then conducted using Kraken2 v2.17.1 with option –use-names^181^. A binary matrix of taxon ID presence and absence in each sample was created and used as input into the UMAP function of umap-learn v0.5.9 with metric jaccard and parameters n_neighbors=50, min_dist=0.1^169^. The embeddings of the UMAPs were colored by the Lineage 3 MGnify biome of the samples and plotted using Datashader v0.18.2^82,170^ [8].

The number of taxa shared between the synthetic metagenome and BaseData digestive system samples was identified using Kraken2 classifications of all full-length contigs, independent of the length-based filtering applied to create the UMAP embedding.

K-mer frequencies (k=4-6) were calculated using Jellyfish v2.3.1 for the synthetic metagenomic contigs and all contigs 7-10kbp in length for the BaseData sample used to fine-tune EDEN, to verify the sequences of the synthetic contigs diverged from the contigs in the fine-tuned sample^168^. A UMAP of Euclidean distances for the k-mer frequencies was then calculated using the UMAP function of umap-learn v0.5.9 with options n_neighbors=20, n_epochs=1000 and min_dist=0^169^. The embeddings of the UMAP were colored by the Lineage 3 MGnify biome of the source sample and plotted using Datashader v0.18.2^169,170^.

### Distant ORF sequence analysis

Generated sequences used pyrodigal (3.0.0) to predict genes^172^. Sequences were length filtered between 250-1500aa. Further filtering was made using hard masking results from tantan default parameters (version 51)^177^. Candidates were then aligned against BaseData, NCBI (2024-02-07), UniProtKB/SwissProt (Release 2025_04 of 08-Oct-2025) and MGnify (v2024_04) using Diamond Blast (v2.1.6) with minimum percentID of 10% and coverage of 0.7^60,82,149,189^. ESMFold (3B V1) was used to solve structures with a length shorter than 600aa^66^. TMAlign (20190822) was used to align the structures of the generated sequences against the highest scoring target from each database^190^. Foldseek (9.427df8a) was then used to identify structural homologous folds from the databases af_proteome and af_swissprot v4, cath50 (v4.3.0), esmatlas30 (v01718804519), pdb (240101), Uniprot50 (v4)^51,60,66,191,192^. MGnify and NCBI peptide records were viewed on December 3, 2025^82,149^.

## Supplementary Information

**Supplementary Figure 1:**
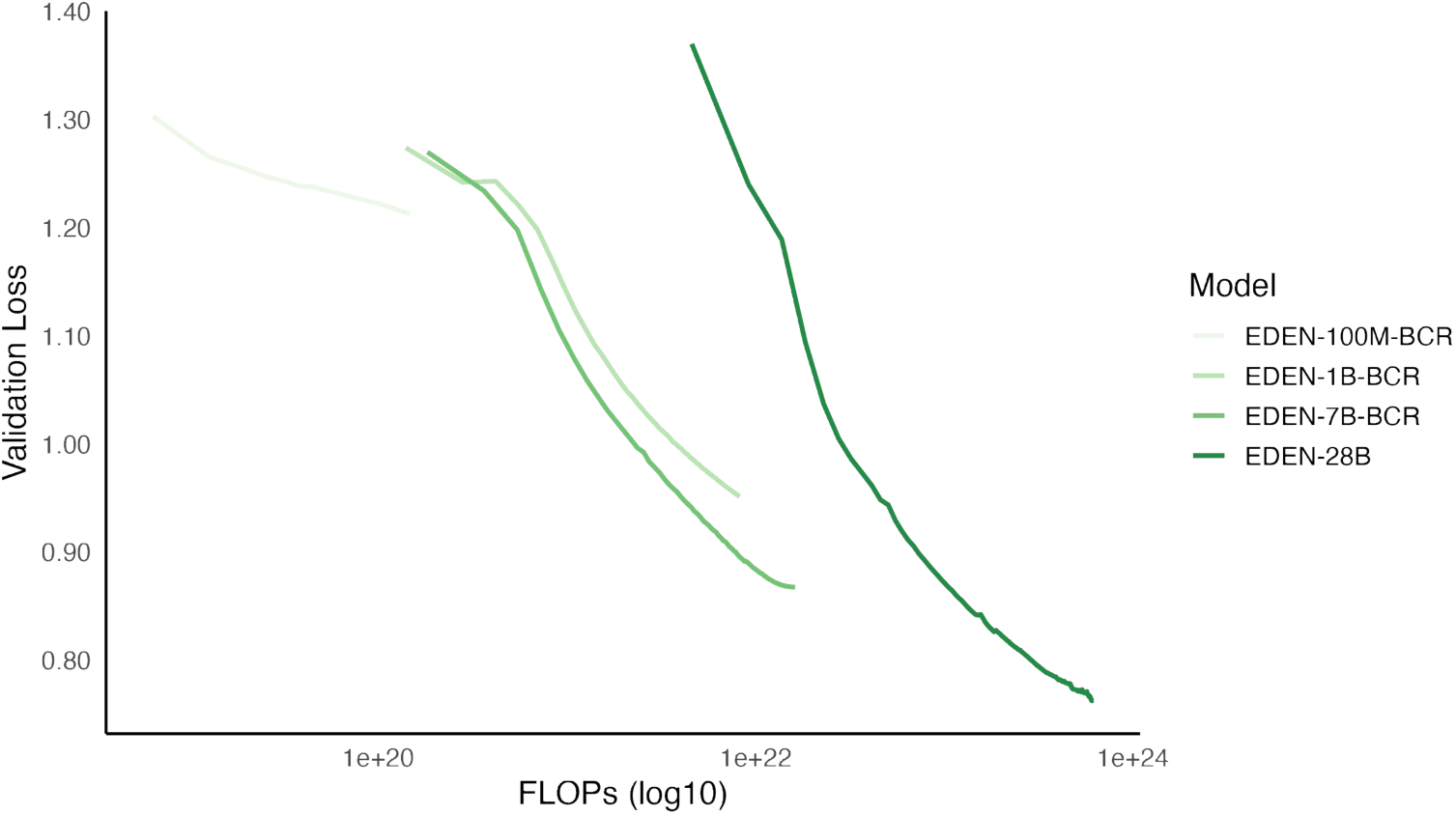
We’re showing scaling behaviour for the EDEN family of models between 100 million and 28 billion parameters. EDEN-100M, EDEN-1B, and EDEN-7B were trained on 350 billion tokens from BaseData. EDEN-28B was trained on 9.7 trillion tokens from BaseData.The diagram displays FLOPS vs validation loss for the various models.

**Supplementary Figure 2:**
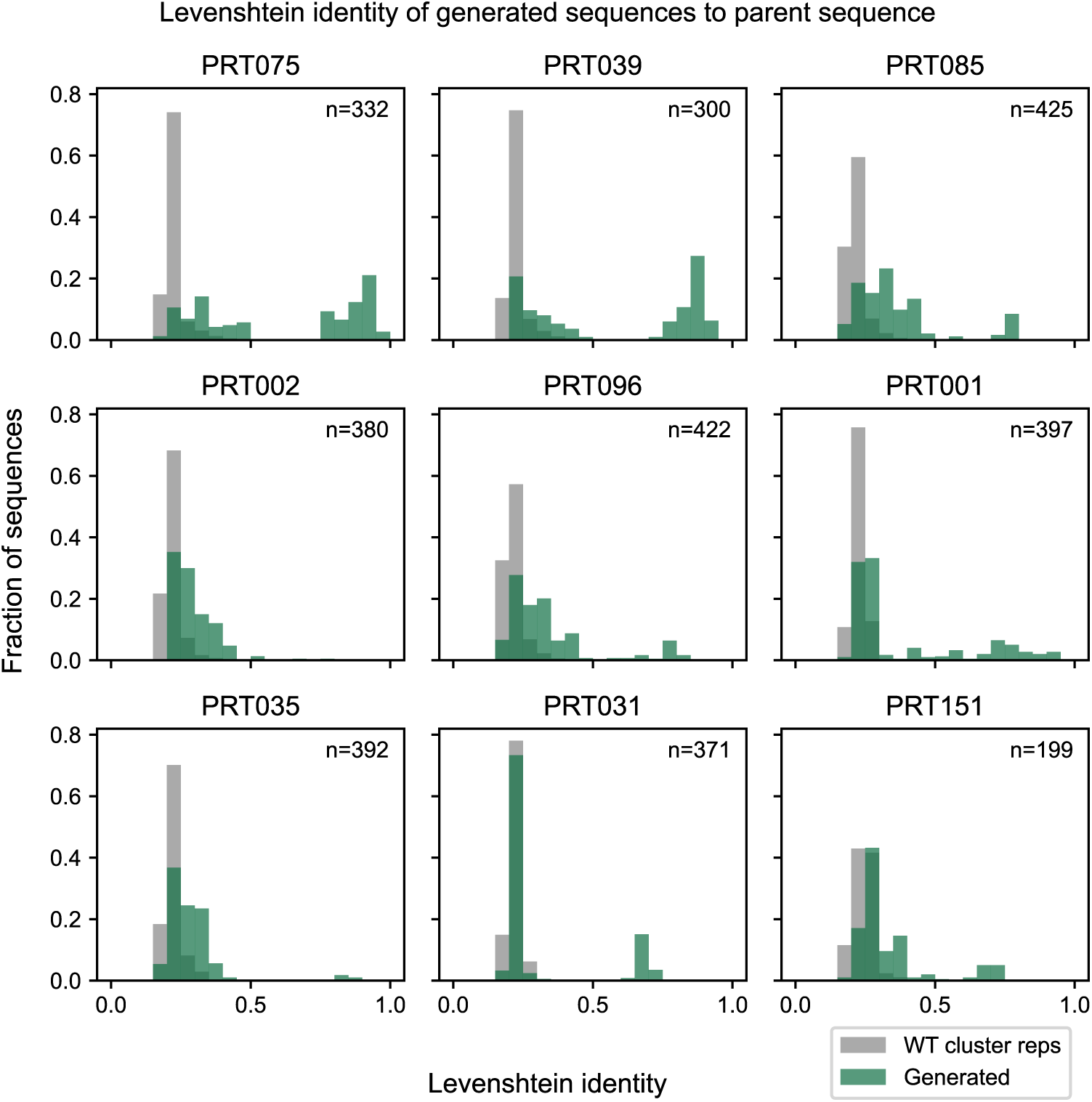
Leventhstein identity of generated sequences vs wild-type LSR from prompt. Distribution of sequence similarity between LSRs. Histograms show the distribution of Levenshtein identity between LSR sequences generated from wild-type prompts and their corresponding parent (prompting) wild-type LSRs (green), and between each parent wild-type LSR and all wild-type LSR cluster representatives (gray; n = 24,330), providing a comparison between model-generated variation and natural sequence diversity. Levenshtein identity = 1 - distance(seq1, seq2) / max_len(seq1, seq2).

**Supplementary Figure 3:**
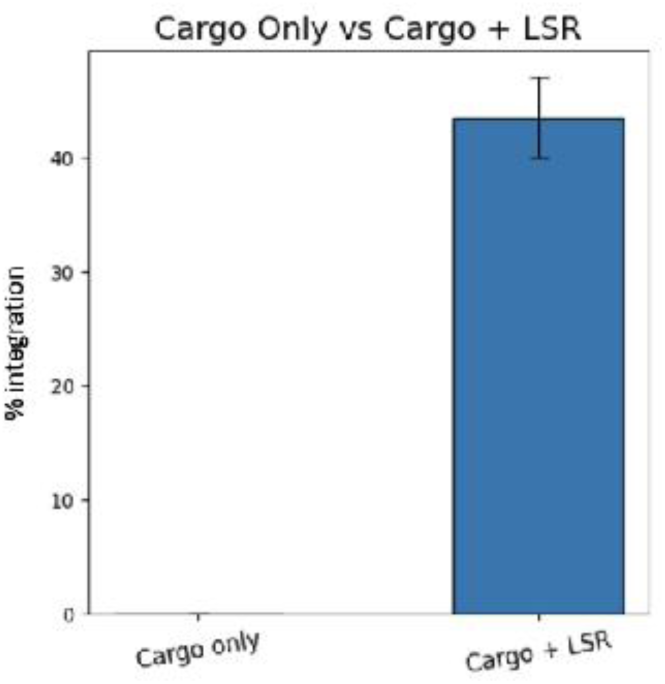
Integration activity of PRT075 in K562 cells. Human K562 cells were transfected with mRNA expressing WT PRT075 along with a plasmid DNA template containing the corresponding att sequence. At 5 days post transfection, DNA was harvested and integration at Safe Harbor 1 - chr7 pseudo site was quantified using dPCR.

**Supplementary Figure 4:**
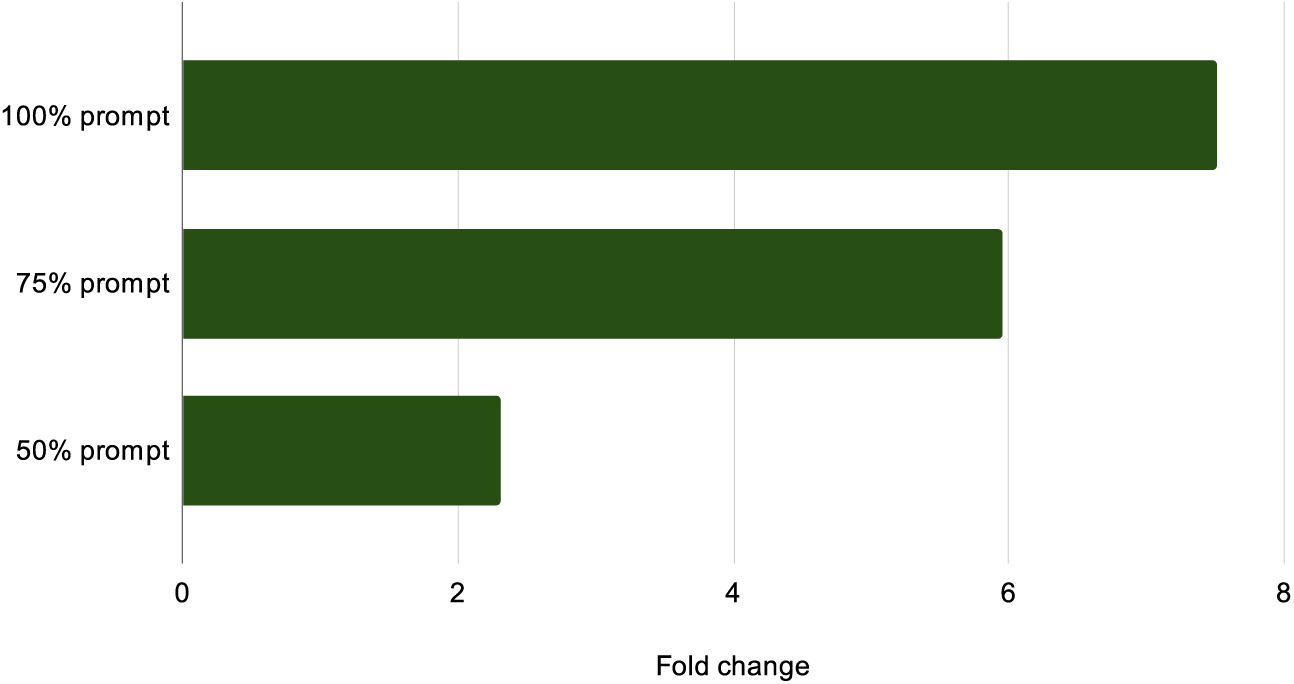
Fold change in the generation rate of EDEN-BR relative to EDEN-28B (calculated as EDEN-BR / EDEN-28B).

**Supplementary Figure 5:**
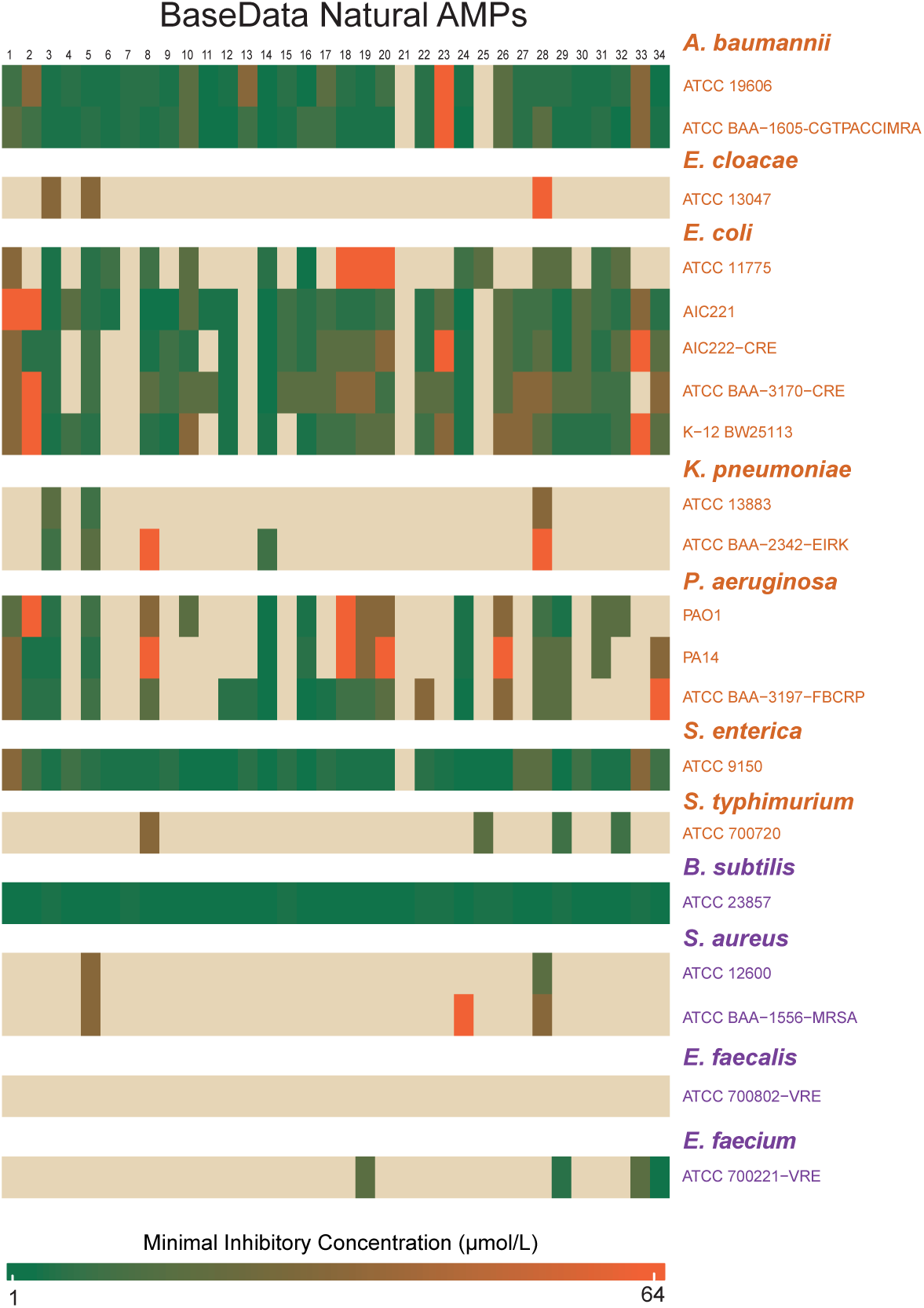
Heatmap showing the results of activity validation assays confirming antimicrobial activity of BaseData peptides against 16 clinically bacterial strains.

**Supplementary Figure 6:**
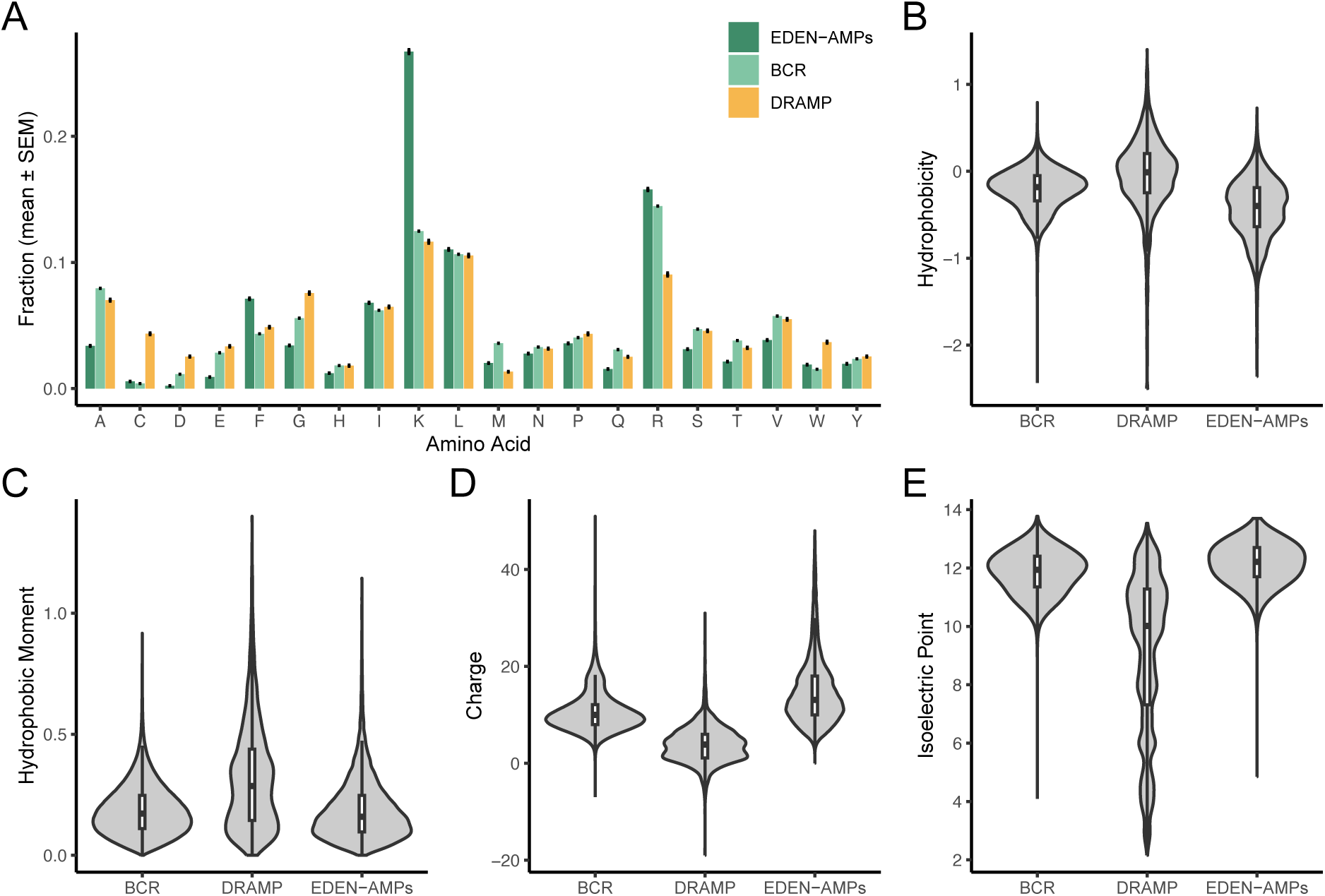
The property distribution of EDEN generated AMPs and natural AMPs from BaseData and DRAMP databases for **A** amino acid composition **B** hydrophobicity **C** hydrophobic moment **D** charge **E** isoelectric point.

## Notes

### Summary of Updates

Correction of 6 spelling errors but no other material changes.

